# Metabolic adaptation to TCA cycle deficiency in a strictly aerobic bacterium

**DOI:** 10.64898/2026.09.17.752348

**Authors:** Changshuo Liu, Eemeli Toivanen, Suvi Santala, Jin Luo, Ville Santala

## Abstract

The tricarboxylic acid (TCA) cycle is central to cellular metabolism, yet its disruption offers an attractive opportunity to redirect carbon toward biotechnological products. A key challenge is that severe TCA cycle interruption often results in auxotrophic phenotypes, limiting their practical application. Here, we used adaptive laboratory evolution to restore growth of the strictly aerobic bacterium *Acinetobacter baylyi* ADP1 following major disruption of the TCA cycle during growth on glucose minimal medium. Evolved strains recovered rapid growth through extensive reorganization of central metabolism, including reduced flux through the oxidative branch of the TCA cycle, enhanced anaplerotic cycling, increased acetate overflow, and remodeling of oxidative phosphorylation. Adaptation involved coordinated changes in quinone-linked electron transport and intracellular redox balancing, enabling continued growth despite reduced respiratory efficiency. These results demonstrate that adaptive evolution can overcome fundamental physiological constraints imposed by TCA cycle disruption and establish a framework for engineering carbon-partitioning phenotypes in aerobic production hosts.

## INTRODUCTION

The tricarboxylic acid (TCA) cycle serves as a central hub of aerobic metabolism. Under aerobic conditions, it oxidizes acetyl-CoA to generate reducing equivalents that drive oxidative phosphorylation, while providing key intermediates for biosynthesis and cellular growth (*1*, *2*). Despite its metabolic robustness and flexibility, its inherent decarboxylation reactions lead to carbon loss as CO_2_, which can limit carbon yield and overall efficiency in microbial bioproduction. This creates a fundamental trade-off between energy generation and carbon conservation. Consequently, the ability to strategically reconfigure or interrupt the TCA cycle has emerged as an attractive objective in metabolic engineering, particularly for improving carbon conservation and enabling efficient biosynthesis of value-added compounds.

Recent studies have demonstrated that metabolic engineering of the TCA cycle can improve product yields and carbon efficiency, establishing TCA manipulation as a powerful strategy in metabolic engineering (*3–5*). However, most TCA-centered studies emphasize reconfiguring or redistributing TCA-associated fluxes rather than on direct interruption of the cycle. For example, in the strictly aerobic yeast *Yarrowia lipolytica*, succinic acid production via reductive TCA engineering requires preservation of overall TCA function to sustain redox balance and cellular growth (*4*). By contrast, interruption of the TCA cycle has been examined in relatively few studies, and primarily in facultatively anaerobic model organisms such as *Escherichia coli* (*6*). Even in this metabolically flexible host, disruption of the TCA cycle has been shown to compromise growth on glucose minimal medium (*6, 7–9*), underscoring the importance of TCA-associated flux for central carbon metabolism. In strictly aerobic bacteria, which lack fermentative alternatives and operate under obligate oxidative metabolism, the TCA cycle plays an even more central role in carbon assimilation and energy generation. Whether TCA cycle manipulation of such rigid oxidative systems can be used not only to improve product yield, but also to direct carbon partitioning between growth and product formation, remains largely unexplored.

*Acinetobacter baylyi* ADP1 (hereafter ADP1) exemplifies this strictly aerobic metabolic lifestyle. It has emerged as a potential lignocellulosic valorization chassis owing to its native capacity to degrade lignin-derived aromatic compounds (*10*, *11*), its intrinsic detoxification of furan derivatives present in lignocellulosic hydrolysates (*12*), and the high tolerance toward hydrolysate components that can be achieved through engineering (*13–15*). As a potential chassis for lignocellulosic valorization, a central limitation of ADP1 lies in its native sugar utilization capabilities. Wild-type ADP1 supports growth only on glucose and cannot metabolize other sugars commonly present in lignocellulosic hydrolysates. This constraint has been partly addressed in previous studies through metabolic engineering, which enabled ADP1 to utilize pentose sugars, namely arabinose and xylose (*16*). In the case of xylose, the engineered strain supported rapid growth (*μ* = 0.73 h⁻¹), corresponding to the highest growth rate reported at the time among non-native microbial hosts engineered for pentose utilization (*16*). Despite this advance, pentose-derived carbon is funneled into central metabolism via α-ketoglutarate (AKG), where continued TCA cycle activity results in substantial diversion of carbon toward biomass formation rather than product synthesis.

In parallel, the previous study investigated whether targeted interruption of the TCA cycle can be used to enforce the conversion of pentose-derived carbon toward AKG-derived product formation in engineered ADP1 (*16*). By deleting *sucAB* to block the canonical TCA cycle and *gabT* to eliminate the γ-aminobutyric acid (GABA) shunt, AKG consumption for biomass formation was effectively prevented, thereby redirecting xylose-derived carbon toward product synthesis. However, simultaneous disruption of the TCA cycle and its major auxiliary sink resulted in severely impaired growth. These observations highlight a fundamental bottleneck in strictly aerobic metabolism, in which restricting TCA flux to preserve carbon for production directly conflicts with the metabolic requirements for growth.

To resolve these constraints, here, we applied adaptive laboratory evolution not merely to recover growth, but to elucidate the metabolic compensatory strategies that emerge when a strictly aerobic network is deprived of its principal carbon-oxidizing path and limited AKG-consumption. Evolutionary adaptation partially restored growth on glucose minimal medium, indicating the existence of compensatory metabolic states that support viability under impaired TCA function. To further characterize these adaptations, ^13^C-based metabolic flux analysis indicated a redistribution of carbon flux within central metabolism. In parallel, transcriptomic analysis revealed system-level regulatory adjustments associated with TCA disruption and growth recovery, providing additional insights into the cellular strategies that accommodate perturbed central carbon metabolism. Together, our results establish TCA interruption as a viable framework for enforcing carbon partitioning in a strictly aerobic bacterium and highlight the potential of the evolved strains as a foundation for future development of AKG-derived bioproduction platforms.

## RESULTS

### Adaptive Laboratory Evolution Restores Growth of a TCA-cycle-deficient Strain

In our previous study, *Acinetobacter baylyi* ADP1 was engineered to utilize xylose and arabinose via the Weimberg pathway, enabling valorization of the hemicellulose fraction of lignocellulosic biomass (*16*). This pathway converts pentoses into the TCA-cycle intermediate AKG, a precursor for numerous valuable compounds (*17*, *18*). However, the high flux through the TCA cycle in this strictly aerobic bacterium competes with product synthesis for AKG. To alleviate this competition, we previously blocked the TCA cycle by deleting *sucAB* and *gabT*, generating strain ASA555 (*16*). The design intended that all pentose-derived carbon would be directed toward the target product via AKG, while glucose would support biomass accumulation. However, disrupting the TCA cycle completely abolished the strain’s ability to grow on glucose as the sole carbon source. To restore growth, we employed adaptive laboratory evolution (ALE).

To initiate ALE, an alternative growth-supporting substrate was required. ASA555 grew well on succinate; thus, we first evolved the strain in mineral salts medium (MSM) containing succinate and glucose. After ∼49 days (∼280 generations), neither of the two independent evolution lines had acquired the ability to grow on glucose (Figure S1). We next tested the growth of ASA555 on rich media, including LB, MSM supplemented with yeast extract, and MSM with casamino acids. Among these, MSM with yeast extract supported the best growth, though much weaker than growth on MSM with succinate. Flux balance analysis (FBA) predicted that the TCA-cycle-deficient strain could achieve a substantial biomass yield on glucose by routing flux through the glyoxylate shunt (Figure S2). We therefore initiated another evolution in MSM with glucose, yeast extract, and acetate to potentially activate this shunt (*19*). Interestingly, the addition of acetate impaired growth compared to medium containing only yeast extract and glucose. Nevertheless, we propagated the populations for ∼51 days (∼200 generations); however, the populations still failed to grow on glucose alone (Figure S1).

We then established two new evolution lines in MSM containing only yeast extract and glucose (Figure S1). Initially, yeast extract was provided at a high concentration (20 g/L) and glucose at a low concentration (4 g/L). As cell optical densities (OD) improved over the course of evolution, yeast extract was gradually reduced while glucose was increased (Figure 1A). After 17 days, one line exhibited marginal growth on glucose alone—a lower final OD and no growth upon transfer to fresh glucose-only medium. This line was subsequently split into two parallel lines. By approximately day 50, both descendant populations had acquired robust growth on glucose as the sole carbon source, retaining this ability after multiple transfers in glucose-only medium. Each of these two lines was then further split (yielding four lines total) and maintained solely on glucose. In contrast, the second original line with glucose and yeast extract still showed no growth on glucose after 50 days of evolution (Figure S1). For the four lines with robust glucose growth, evolution was continued until days 99–100. During this phase, the glucose concentration was lowered from 6 g/L to 2.8 g/L to prevent medium acidification caused by rapid oxidation of glucose to gluconate. Notably, this reduction did not decrease the final OD of the cultures. Six individual clones were isolated from the endpoint populations of the four evolution lines (Figure 1A) and designated ASA557-1 to ASA557-6. All evolved isolates grew on glucose and displayed similar growth profiles (Figure 1B). Interestingly, the evolved isolates were nearly unable to grow on succinate as the sole carbon source.

**Figure 1.**
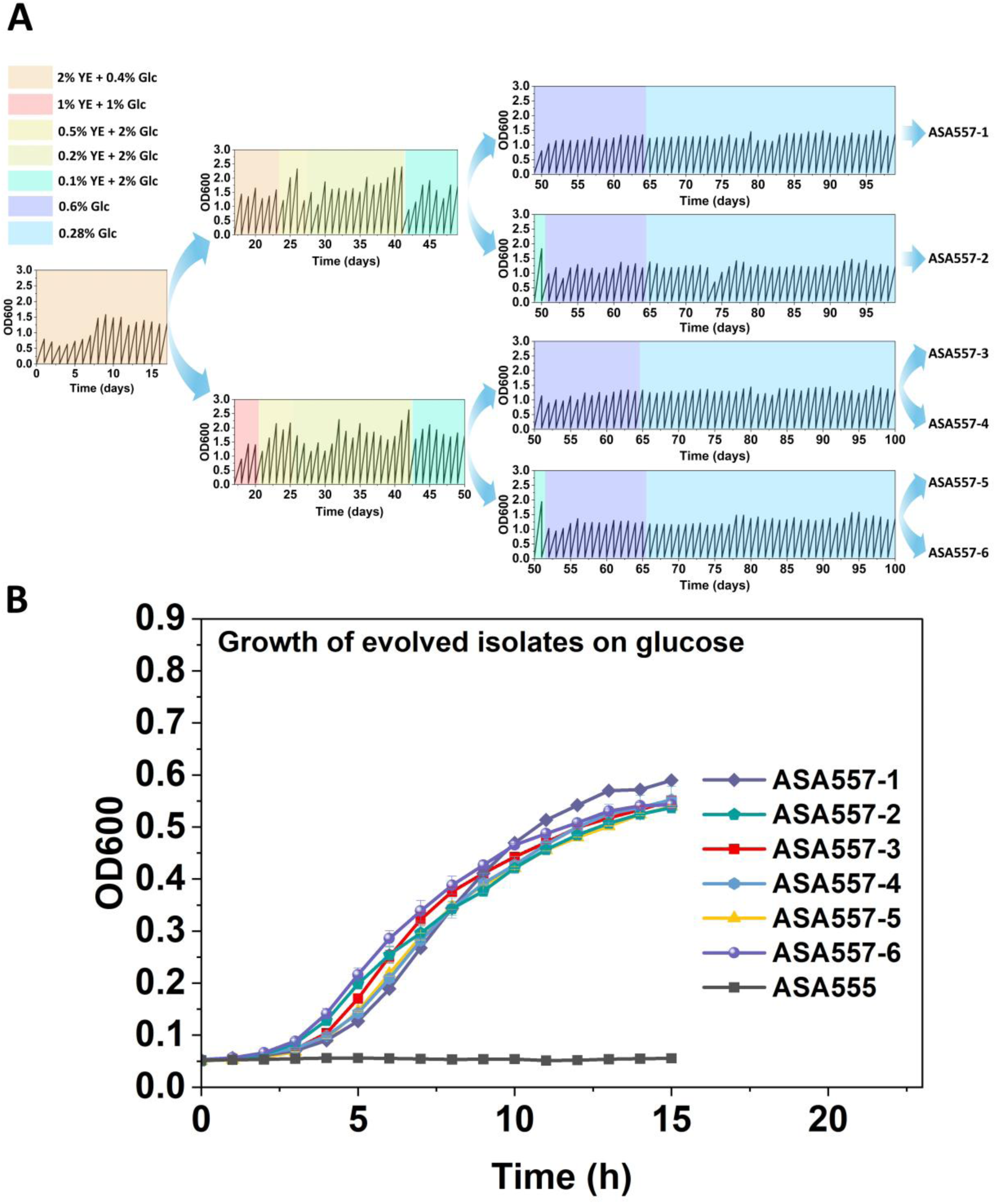
Adaptive laboratory evolution restores growth of the TCA-cycle deficient strain ASA555 (Δ*sucAB*, Δ*gabT*::*tdk*/*kan*) on glucose as the sole carbon source. (**A**) Evolution trajectories of ASA555. Cells were first evolved in mineral salts medium supplemented with yeast extract and glucose (YE + Glc), followed by passage in glucose alone, yielding multiple evolved isolates. The percentage symbol (%) in the legends denotes weight/volume (w/v). (**B**) Growth comparison of evolved isolates and the parental ASA555 strain on glucose as the sole carbon source. Data represent mean values and error bars indicate standard deviations from two independent biological experiments. Abbreviations: YE, yeast extract; Glc, glucose.

### Genetic Mutations Associated with Evolution

To identify mutations underlying the restored growth on glucose, we performed whole-genome sequencing of the six evolved isolates and the parental strain ASA555. Mutations unique to the evolved isolates are listed in Table 1.

**Table 1.** Mutations identified in the evolved isolates.

| Related gene * | Position * | Description | DNA change | Effect | ASA 557-1 | ASA 557-2 | ASA 557-3 | ASA 557-4 | ASA 557-5 | ASA 557-6 |
| --- | --- | --- | --- | --- | --- | --- | --- | --- | --- | --- |
| *gyrB* | 4226 | DNA gyrase subunit | C→T | R34C (CGT→TGT) | + |  |  |  |  |  |
| *rpoC* | 308764 | DNA-directed RNA polymerase beta' | G→T | G278C (GGT→TGT) |  |  |  |  | + | + |
| *tRNA<sup>Lys</sup>_16* | 435023-435205 | Transfer RNA of lysin | Δ182 bp | Large deletion |  | + |  |  |  |  |
| *ACIAD0801* | 793329 | Putative membrane protein | C→T | M668I (ATG→ATA) | + | + | + | + | + | + |
| *ACIAD0955*, *ACIAD0956* (CDS 29-270/270) | 943890-944967 | Hypothetical proteins of the Tn5613 transposon | Δ1078 bp | Transposon mediated large deletion |  |  |  |  | + |  |
| From *pbpA* (CDS 1-1182/2028) to *ACIAD1847* (CDS 1-841/1164) | 1095113-1852976 | NA | 757864 bp inversion | Inversion |  | + |  |  |  |  |
| *ACIAD1407* → / → *hupB* | 1404756 | Putative poly(hydroxyalkanoate) granule associated protein/DNA-binding protein HU-beta | G→A | Intergenic (+69/-95) | + | + | + | + | + | + |
| *ACIAD2017* ← / → *acoD* | 2014075-2014076 | Putative transcriptional regulator/Acetaldehyde dehydrogenase 2 | +IS insertion, +GAG | Intergenic (-76/-85) |  |  |  | + |  |  |
| *glcB* (CDS 1-173/2163), *ACIAD2336* (CDS 689-1143/1143) | 2303831-2304808 | Malate synthase G, putative ATPase | Δ979 bp | Large deletion |  |  | + | + | + | + |
| *ACIAD2336* | 2305253 | Putative ATPase | C→G | G82R (GGT→CGT) | + | + |  |  |  |  |
| *rpsF* | 2400138 | 30S ribosomal protein S6 | A→C | Q17H (CAA→CAC) |  |  |  | + | + | + |
| *rpsF* | 2400304 | 30S ribosomal protein S6 | G→A | E73K (GAA→AAA) |  |  | + |  |  |  |
| *greA* (CDS 383-477/477), *ACIAD2863* (CDS 1-91/459) | 2802807-2803135 | Transcription elongation factor, putative universal stress protein A | Δ329 bp | Large deletion |  |  | + |  |  |  |
| *sdhA* | 2822114 | Succinate dehydrogenase subunit | A→T | S300T (TCA→ACA) | + | + | + | + | + | + |
| *sdhC* ← / → *gltA* | 2824820 | Succinate dehydrogenase Subunit/citrate synthase | G→A | Intergenic (-1030/-13) | + | + | + | + | + | + |
| *rpoD* | 2854342 | RNA polymerase sigma factor | C→T | R102C (CGC→TGC) | + |  |  |  |  |  |
| From *ACIAD2956* (CDS 1-546/906) to *qseC* (CDS 976-1326/1326) | 2886266-2889687 | NA | Δ3,422 bp | Large deletion | + | + |  |  |  |  |
| *qseC* | 2890008 | Two-component sensor kinase transcription regulator | G→A | Q219* (CAG→TAG) |  |  | + | + | + | + |
| *rho* | 2967670 | Transcription termination factor | G→C | G61R (GGT→CGT) | + | + |  |  |  |  |

\* Locus IDs and mutation positions were assigned according to the ASA555 genome (Data S5), which was derived from the reference genome CR543861 (GenBank entry). Locus IDs are provided when gene names are unavailable.

Six loci were mutated in all six isolates: *ACIAD0801*, the intergenic region between *ACIAD1407* and *hupB*, *ACIAD2336*, *sdhA*, 13 bp upstream of *gltA*, and *qseC*. Given that ACIAD0801 is annotated as an integral membrane transporter, the M668I substitution is unlikely to directly contribute to the restored growth on glucose. The substitution in the *ACIAD1407*–*hupB* intergenic region may have impact on *hupB* expression but it does not affect the adjacent coding sequences. The remaining four mutations are more likely to be functionally significant. Two are connected to the TCA cycle: (1) the *sdhA* mutation replaces serine with threonine (S300T) in succinate dehydrogenase subunit A, a position close to the predicted active-site residue (position 296) that acts as a proton acceptor, and (2) a G to A transition 13 bp upstream of the *gltA* start codon, which may alter transcription of the citrate synthase. Both mutations are identical across all six isolates, suggesting they arose before the divergence of the evolution lines, at a stage when only one line was present (Figure 1A).

Of the two remaining significant mutations *ACIAD2336* encodes a cell division ATPase. In isolates ASA557-1 and ASA557-2, it carries a G82R mutation—a substitution of a small, neutral glycine with a large, positively charged arginine—which likely disrupts protein function. In ASA557-3 to ASA557-6, the same gene contains an in-frame deletion (nucleotides 689–1143) that presumably results in complete loss of function. The *qseC* gene, which encodes a two-component sensor histidine kinase, is also mutated in all six isolates: a Q219* nonsense mutation truncates the protein in ASA557-3 to ASA557-6, whereas ASA557-1 and ASA557-2 carry an in-frame deletion (nucleotides 976–1326). The identical mutations in *ACIAD2336* and *qseC* shared by ASA557-1 and ASA557-2, and the distinct but phenotypically similar mutations in ASA557-3 to ASA557-6, indicate that these mutations appeared after the initial split of one evolution line into two, but before the two lines were split further. The convergent inactivation of the same genes through independent evolution lines strongly implicates that both *ACIAD2336* and *qseC* are important in the adaptive phenotype.

Unexpectedly, a loss-of-function mutation occurred in *glcB* (encoding malate synthase, the second enzyme of the glyoxylate shunt) in isolates ASA557-3 to ASA557-6, suggesting that the glyoxylate shunt is blocked in these strains. This finding, combined with the FBA prediction that the TCA-cycle-deficient strain could grow via the glyoxylate shunt, indicates that the evolved isolates have rewired central metabolism through alternative routes. Additionally, besides *qseC*, all isolates harbored mutations in other genes with potentially global regulatory effects: *gyrB* in ASA557-1, *rpoC* in ASA557-5 and ASA557-6, *rpsF* in ASA557-3 to ASA557-6, and *rho* in ASA557-1 and ASA557-2. These mutations may further modulate transcription, translation, or DNA topology, contributing to the evolved growth phenotype.

### Reverse Engineering Confirms the Adaptive Mutations

To validate the key mutations responsible for restored growth on glucose, we performed reverse engineering. We initially attempted a transformation-based enrichment strategy previously described for *Acinetobacter baylyi* (*20*). Specifically, we PCR-amplified each mutated locus from strain ASA557-6. The strain was chosen because it carries fewer mutations than the other evolved isolates, including the *rpoC*, *ACIAD0801*, *ACIAD1407*-*hupB* intergenic, *glcB*-*ACIAD2336*, *rpsF*, *sdhA*, *gltA* upstream, and *qseC* mutations. The parental strain ASA555 was grown in MSM supplemented with yeast extract and glucose, and the amplified DNA fragments were added to the culture either individually or as a mixture. DNA was added at each serial passage to facilitate integration, exploiting ADP1’s natural competence and high recombination efficiency. After multiple transfers, the populations were tested for growth on glucose alone. No culture acquired the ability to grow on glucose as the sole carbon source. This result indicates that single mutation is not sufficient, and that multiple mutations must be simultaneously present in the same cell to produce a selectable growth phenotype. While the transformation mixture approach may integrate several fragments, its efficiency decreases as the number of required integrations increases, and it likely yields cells carrying different, incomplete subsets of mutations. Consequently, cells that acquired the full set of mutations were too rare to be enriched under the selective pressure.

We therefore introduced the mutations sequentially, focusing on those shared by all six evolved isolates: *sdhA* (S300T), the G to A substitution 13 bp upstream of *gltA*, *qseC* (Q219*), and *ACIAD2336* deletion (Figure 2A). First, the *sdhA* and *gltA* upstream mutations were introduced together due to their adjacent loci into the parental strain to generate strain RE1. RE1 achieved a higher final optical density than ASA555 in MSM with yeast extract and glucose and exhibited very slight growth on glucose as the sole carbon source. Next, the *qseC* (Q219*) mutation was introduced into RE1, yielding RE2. However, RE2 did not show substantially improved growth on glucose relative to RE1 (Figure 2B). In ASA557-6, both *glcB* and *ACIAD2336* contain loss-of-function mutations; because *glcB* remains intact in some evolved isolates, we specifically knocked out only *ACIAD2336* in the RE2 background, creating strain RE3. Comparison between the engineered strains and the parental strain revealed that the first three mutations produced minimal additive effect; a marked improvement occurred only upon introduction of the fourth mutation, the *ACIAD2336* knockout (Figure 2B). RE3, which carries all four common mutations, largely regained the ability to grow on glucose, but its biomass yield and growth remained lower than those of ASA557-6. Thus, the shared mutations in *sdhA*, *gltA*, *qseC*, and *ACIAD2336* account for the adaptive phenotype to some extent, but full restoration requires additional strain-specific mutations, likely including those with global regulatory effects.

**Figure 2.**
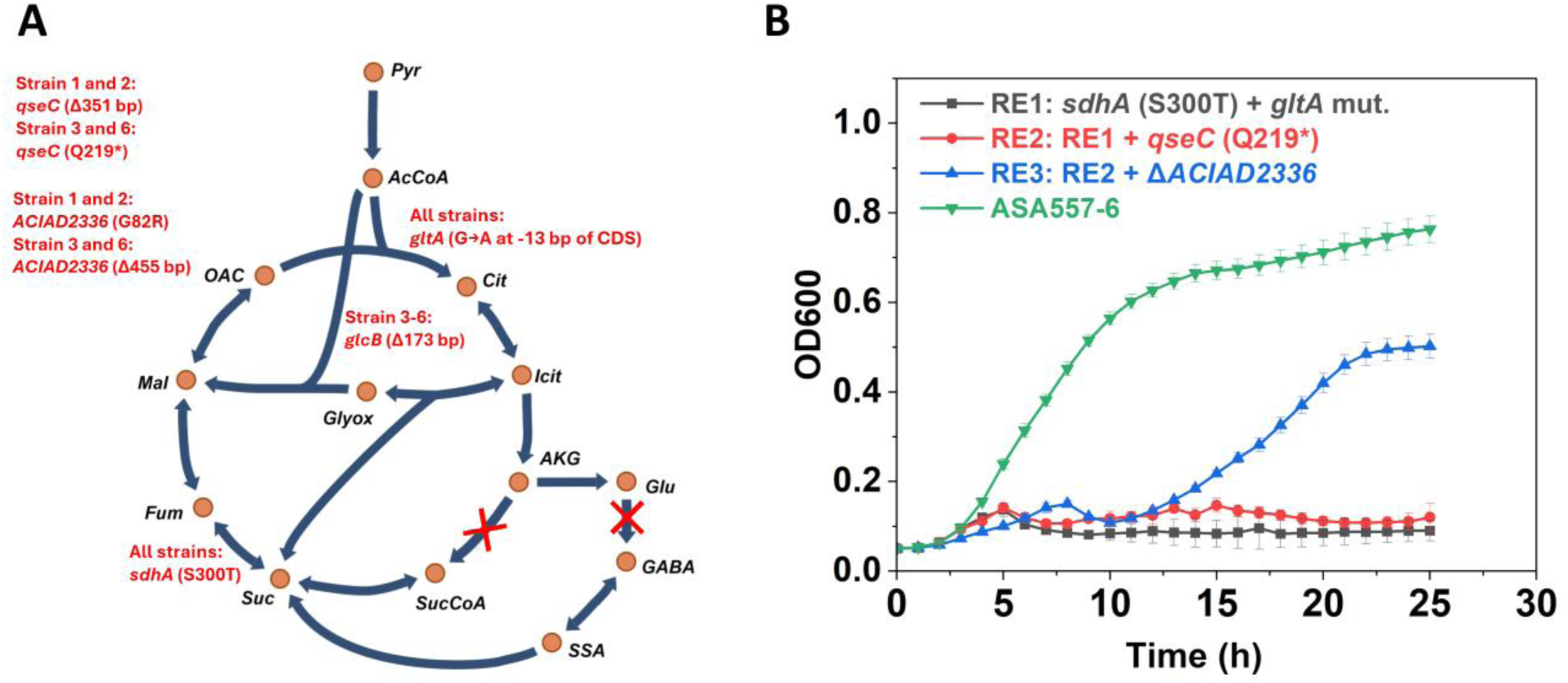
Key mutations validated by reverse engineering. (**A**) Key mutations found to be important for growth recovery on glucose. Several mutations are associated with the TCA cycle. Red crosses indicate pathways blocked in the parental strain ASA555. (**B**) Growth comparison of reverse-engineered strains (RE1, RE2, and RE3) and the evolved isolate ASA557-6 in mineral salts medium supplemented with glucose as the sole carbon source. Data represent mean values and error bars indicate standard deviations from three independent biological experiments. Abbreviations: Pyr, pyruvate; AcCoA, acetyl-CoA; OAC, oxaloacetate; Cit, citrate; iCit, isocitrate; AKG, α-ketoglutarate; SucCoA, succinyl-CoA; Suc, succinate; Fum, fumarate; Mal, malate; Glyox, glyoxylate; Glu, glutamate; GABA, γ-aminobutyric acid; SSA, succinic semialdehyde.

### Metabolic Rewiring in the TCA-cycle-deficient Strain

Although the TCA-cycle-deficient strain was evolved to grow on glucose, the evolved isolates still differ phenotypically from the strain with an intact TCA cycle. At equivalent glucose concentrations, the evolved strains reached a lower final optical density. For example, when cultivated in MSM with 10 mM (1.8 g/L) glucose as the sole carbon source, the reference strain ASA549 reached an OD600 above 2, whereas the evolved isolates ASA557-3 and ASA557-6 reached an OD600 of approximately 1 (Figure S3). However, the evolved strains exhibited a higher growth rate during the log phase, as indicated by steeper slopes in the logarithmic scale growth curves. Unlike the reference strain, increasing glucose concentration had little effect on the final OD of the evolved isolates. This limitation may stem from the fact that, in the evolved strains, the first step of the Entner-Doudoroff (ED) pathway, the oxidation of sugar to its sugar acid, is coupled to growth.

We further examined the growth and substrate consumption of ASA557-3 in a fed-batch bioreactor. Cells were cultivated in MSM containing 44.4 mM (8 g/L) glucose. After 17 h, glucose was fully oxidized to gluconate, and the dissolved oxygen (DO) rose sharply despite a large amount of gluconate remaining in the medium (Figure S4). At this point, xylose was added in pulses. Xylose metabolism proceeds through the heterologous Weimberg pathway, and its first step—oxidation to xylonate—is catalyzed by the same enzyme that oxidizes glucose (*16*). Each xylose addition was accompanied by a drop in DO and an increase in biomass, as reflected by OD600 and real-time backscatter (Figure S4A). As xylose was fully oxidized, DO increased again and biomass accumulation plateaued, which supports the coupling of the oxidation step to growth. From 17 h onward, the culture showed slow consumption of gluconate and accumulation of xylonate, acetate, and AKG (Figure S4A). A similar phenomenon was also observed for ASA557-6 under a similar experiment setup. We next performed a similar experiment with ASA557-6 and supplemented formic acid, hypothesizing that its oxidation to CO2 would support continued growth and consumption of the remaining substrates. As expected, formic acid supplementation enabled the cells to continue growing and consuming the remaining substrates (Figure S4B). Similar effect on growth was observed on RE3 (data not shown).

To investigate the metabolic rewiring underlying the evolved phenotype, we quantified intracellular metabolite levels and performed ^13^C metabolic flux analysis on the evolved isolate ASA557-6 and the reference strain ASA549. For both experiments, cells were cultivated in MSM with 11.1 mM (2 g/L) glucose as the sole carbon source and 50 mM MOPS for pH buffering. Samples for metabolite quantification were taken at OD600 ≈ 0.55 (log phase for both strains) and OD600 ≈ 0.95 (log phase for ASA549; early stationary phase for ASA557-6). Figures 3A and 3B show the relative abundance of selected metabolites in ASA557-6 compared to the reference strain ASA549 at the two sampling points; absolute concentrations are provided in Figure S5. In both strains, glutamate was the most abundant of the measured metabolites (Figure S5). Notably, the abundance of TCA-cycle intermediates and derived metabolites differed markedly between the two strains (Figure 3A and 3B). AKG was approximately fourfold more abundant in ASA557-6 than in ASA549 at OD600 ≈ 0.55, and this difference was even greater at OD600 ≈ 0.95. Glutamate showed a similar trend, being more abundant at both time points. These results are consistent with the *sucAB* deletion, which blocks conversion of AKG to succinyl-CoA. In contrast, citrate was less abundant in ASA557-6, and downstream TCA-cycle intermediates—including succinate, fumarate, and malate—were even more severely depleted, reflecting reduced replenishment due to the cycle blockage. Interestingly, GABA was detected in ASA557-6 but not in ASA549 (Figures S6 and S7), consistent with the *gabT* deletion.

**Figure 3.**
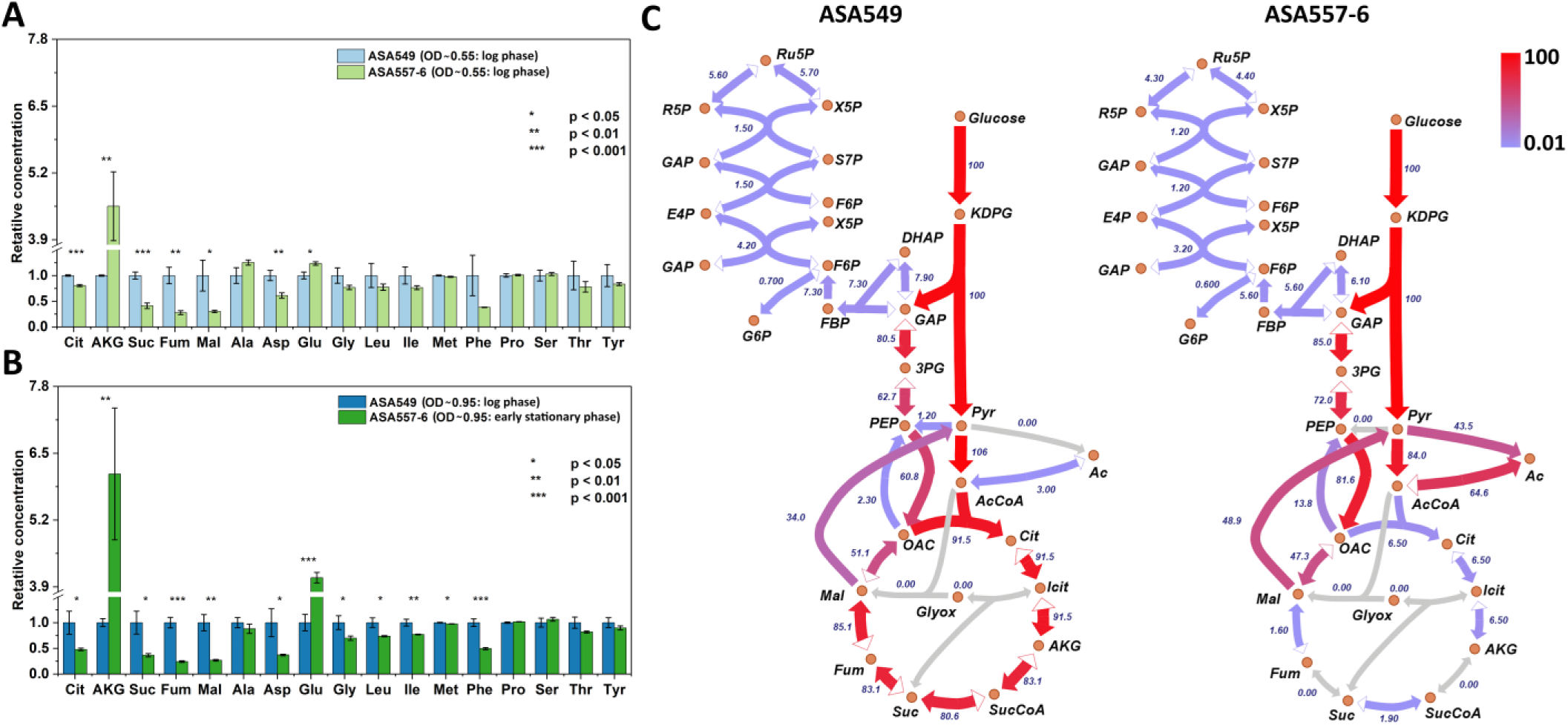
Metabolic rewiring of the TCA-cycle deficient strain. (**A**) Relative intracellular metabolite concentrations in strain ASA557-6 compared to ASA549 at OD600 ≈ 0.55. (**B**) Relative intracellular metabolite concentrations at OD600 ≈ 0.95. For both time points, equivalent biomass samples were harvested for metabolite extraction: (1) OD600 ≈ 0.55 (log phase for both strains) and (2) OD600 ≈ 0.95 (log phase for ASA549; early stationary phase for ASA557-6). Data represent mean values and error bars indicate standard deviations from three independent biological experiments. Statistical significance was determined by Student’s t-test (* p < 0.05, ** p < 0.01, *** p < 0.001). (**C**) Carbon flux maps of ASA549 and ASA557-6 determined by ^13^C metabolic flux analysis using parallel labeling experiments with 50% [U-^13^C] D-glucose and [1-^13^C] D-glucose. Detailed flux values and confidence intervals are provided in Data S2.Abbreviations: Ru5P, ribulose 5-phosphate; R5P, ribose 5-phosphate; X5P, xylulose 5-phosphate, S7P, sedoheptulose 7-phosphate; E4P, erythrose 4-phosphate; F6B, fructose 6-phosphate; FBP, fructose 1,6-bisphosphate; GAP, glyceraldehyde3-phosphate; DHAP, dihydroxyacetone phosphate; 3PG, 3-phosphoglycerate; PEP, phosphoenolpyruvate; Pyr, pyruvate; AcCoA, acetyl-CoA; OAC, oxaloacetate; Cit, citrate; Icit, isocitrate; AKG, α-ketoglutarate; SucCoA, succinyl-CoA; Suc, succinate; Fum, fumarate; Mal, malate; Glyox, glyoxylate.

The flux distributions determined by ^13^C metabolic flux analysis were in agreement with the metabolite quantification data and the genotypes of the engineered strains. Relative to ASA549, ASA557-6 showed markedly reduced flux through the TCA cycle and no flux from AKG to succinyl-CoA (Figure 3C). A substantial fraction of carbon was directed to acetate (Figure 3C); however, whether acetate was produced from pyruvate or acetyl-CoA could not be confidently resolved with the labeling strategy employed (parallel experiments using 50% [U-^13^C]glucose and [1-^13^C]glucose). In both ASA557-6 and ASA549, flux through the glyoxylate shunt was negligible (Figure 3C). A striking feature of the ASA557-6 flux map was a substantial flux from phosphoenolpyruvate (PEP) to oxaloacetate (OAC) (Figure 3C). This was further supported by the conversion between OAC and malate—a reaction well resolved by our labeling setup—showing high net flux from OAC to malate in ASA557-6, whereas the opposite direction was observed in ASA549 (Figure 3C). These patterns are also reflected in the mass isotopomer distributions of proteinogenic amino acid fragments from cultures grown on [1-^13^C]glucose (Data S3 and Figure S8; data for 50% [U-^13^C]glucose are shown in Figure S9, though their interpretation is less intuitive). For example, with [1-^13^C]glucose, a higher fraction of labeled aspartate and glutamate was observed in ASA557-6. This is because the carbon flux entering the TCA cycle via acetyl-CoA loses the ^13^C label as CO_2_, whereas carbon routed from PEP to oxaloacetate retains the label. Detailed flux values and confidence intervals for each reaction are provided in Data S2.

### Transcriptomic Changes in the Evolved Strain

To investigate transcriptional differences between the evolved isolates and the reverse-engineered strain, we performed whole transcriptome sequencing under the same cultivation conditions used for intracellular metabolite quantification. Cultures of evolved isolates ASA557-3 and ASA557-6, reverse engineered strain RE3 and reference strain ASA549 were sampled during exponential growth (OD 600 ∼0.35). Differential expression analysis was performed using ASA549 as the reference strain. Shrunken log two-fold changes and their padj-values for differential expression analysis are available in supplementary Data S4. Principal component analysis showed clear separation between the reference, evolved isolates, and the reverse-engineered strain, with tight clustering of biological replicates (Fig. S10). Separation between reference and modified strains was mainly associated with the presence of *sucAB* and expression levels of central metabolism and carbon catabolism related genes, whereas differentiation between RE3 and evolved isolates was dominated by genes involved in respiratory and redox metabolism. Figure 4 summarizes pathway-level transcriptional responses and representative genes associated with the major expression changes observed in ASA557-3, ASA557-6, and RE3.

**Figure 4.**
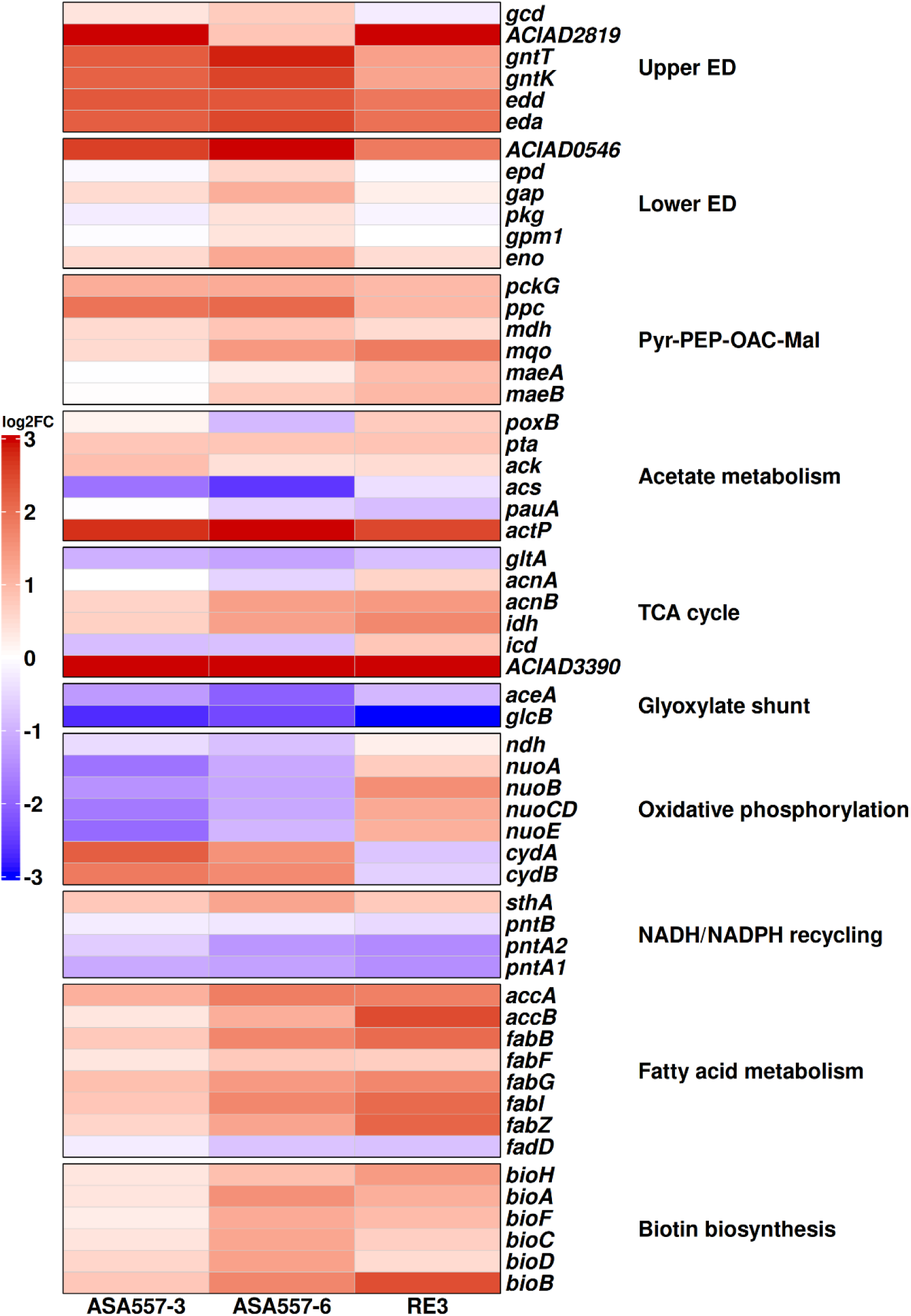
Heatmap of transcription log2 fold changes of representative genes for adapted isolates ASA557-3, ASA557-6 and reverse engineered strain RE3. Abreviations: ED, Entner–Doudoroff pathway; Pyr, pyruvate; PEP, phosphoenolpyruvate; OAC, oxaloacetate; Mal, malate; TCA, tricarboxylic acid cycle.

Expression patterns indicated extensive reorganization of central carbon metabolism following TCA cycle disruption. Despite reduced TCA-cycle fluxes measured by MFA, several genes associated with the oxidative branch of the TCA cycle showed increased expression. In contrast, expression of the primary citrate synthase gene *gltA* was reduced, whereas expression of *prpC*, encoding an enzyme with secondary citrate synthase activity, was increased in the evolved isolates (*21*). Similarly, expression of the alternative isocitrate dehydrogenase *idh* increased in all strains, whereas expression of the canonical enzyme *icd* was reduced in the evolved isolates but increased in RE3. The engineered *sucAB* deletion eliminated expression of the AKG dehydrogenase complex, while ACIAD3390, encoding a putative succinyl-CoA:acetate CoA-transferase/hydrolase, was among the most strongly upregulated genes. Downstream of succinate, fumarate-to-OAC reactions including *fumA*, *mqo*, and *mdh* were generally upregulated, whereas glyoxylate-shunt genes *aceA* and *glcB* were strongly repressed, consistent with the absence of flux through this pathway.

Transcriptomic data also supported redirection of carbon through the Entner-Doudoroff pathway, acetate metabolism, and anaplerotic cycling. Expression of *gcd*, ACIAD2819, *gntT*, *gntK*, *edd* and *eda* were elevated in all strains, indicating enhanced upper glucose catabolism. In contrast lower ED genes *gap, epd, eno* saw stronger upregulation in ASA557-6 while ACIAD0546, a putative NADP-dependent G3P dehydrogenase was heavily upregulated across strains. Genes associated with acetate production (*pta* and *ack*) and transport (*actP*) were modestly upregulated while *poxB* saw slight downregulation in ASA557-6, whereas genes associated with acetate reassimilation were generally reduced. In parallel, strong induction of *ppc* and *pckG* together with increased expression of malic enzyme genes *maeAB* with *mqo* and *mdh* were consistent with enhanced utilization of the PEP-OAC-malate-pyruvate node and in line with the flux distributions observed by MFA.

The most pronounced transcriptional differences between the evolved isolates and RE3 were observed in pathways associated with oxidative phosphorylation, electron transport chain (ETC) and intracellular redox metabolism. Genes of the NADH dehydrogenase I (*nuo*) operon were broadly downregulated in the evolved isolates but upregulated in RE3. Similarly the NADH dehydrogensae *ndh* was downregulated in isolates but remaided unchanged in RE3. In contrast, the cytochrome bd oxidase genes *cydAB* displayed the opposite pattern and were strongly induced in the evolved isolates while repressed in RE3. In contrast the alternative terminal oxidases *cioA/B* were downregulated in all strains. Expression of cytochrome c oxidase (*cyo*) was modestly elevated only in the evolved isolate ASA557-6, whereas ATP synthase genes *atp* were modestly upregulated in both evolved and engineered backgrounds. Additional differences were observed in cofactor-balancing reactions. The membrane-bound transhydrogenase *pnt* was strongly downregulated while the soluble transhydrogenase *sthA* was strongly induced, and expression patterns of *idh*/*icd*, *mqo*/*mdh* and ACIAD0546/*gap-pkg* suggested remodeling of NAD(P)H-associated metabolism.

Genes associated with biosynthesis and growth-related functions were also transcriptionally affected. Fatty-acid biosynthesis was among the most strongly induced pathways, with elevated expression of *accA/B/C/D*, *bccA*, and multiple *fab* genes. Consistent with increased lipid synthesis, biotin biosynthesis genes (*bioA*, *bioB*, and *bioD*) were upregulated, whereas *fadD*, involved in fatty-acid degradation, was repressed. Amino-acid biosynthesis was similarly redistributed toward pathways connected to central metabolism. Biosynthetic routes derived from OAC and 3PG were generally upregulated, whereas branched-chain amino-acid biosynthesis pathways were reduced. Despite high intracellular glutamate levels (Fig. 3A and 3B), expression of several glutamate-associated genes including *gdhA*, *gltBD*, and *glnA* remained unchanged or reduced in the evolved isolates while *gltBD* and *glnA* were upregulated in RE3. In contrast, arginine biosynthesis pathways downstream of glutamate were consistently upregulated and the cyanophycin synthetase CphA showed moderately increased expression. Because both arginine and cyanophycin synthesis consume glutamate-derived nitrogen, their increased expression may represent a response to elevated glutamate availability and provide routes for redistribution or storage of excess nitrogen generated by TCA-cycle disruption.

In addition to metabolic genes, transcriptional changes were observed in several regulatory systems. Although mutations were identified in genes associated with transcription, translation, and chromosome organization, including *rpoC*, *greA*, and *rpsF* (Table 1), these genes generally showed limited changes in transcript abundance. In contrast, the two-component system *qseBC* was consistently downregulated across all strains in addition to the evolved nonsense mutation in *qseC*. The ribosomal protein gene *rpsF* showed modest induction in ASA557-6, whereas the histone-like DNA-binding protein *hupB* was strongly repressed in the evolved isolates but remained near reference levels in RE3. Together, these transcriptional profiles suggest that growth restoration involved coordinated reorganization of central metabolism, respiratory functions, and intracellular redox balancing.

## DISCUSSION

TCA cycle redirection is a promising metabolic engineering strategy which allows carbon flux partitioning away from oxidative metabolism toward desired bioproduction pathways. However, disruption of such a central metabolic pathway can impose severe constraints on precursor supply, redox balancing, and energy generation (*6*, *16*). Here metabolic rewiring during adaptive evolution enabled ADP1 growth on glucose minimal media despite TCA cycle disruption. In ALE-derived isolates, we observed extensive metabolic reorganization, including redistribution of central metabolic fluxes, emergence of acetate overflow, and altered intracellular metabolite pools (Fig. 3). Additionally, pathways involved in oxidative phosphorylation and redox cofactor balancing saw drastic changes (Data S4 and Fig. 4), highlighting the strong dependence of this strictly aerobic bacterium on respiratory metabolism. Together, these adaptations appear to alleviate the metabolic constraints imposed by TCA cycle disruption by supporting precursor supply, cofactor metabolism, and energy conservation. ADP1 provides a particularly useful model for examining these responses because glucose catabolism proceeds primarily through the Entner-Doudoroff pathway, resulting in a less redundant central metabolic network than in organisms such as *E. coli*. Consequently, adaptive responses to TCA-cycle disruption can be interpreted more directly in terms of changes in carbon flux, respiration, and redox balancing. Collectively, these results illustrate the flexibility of bacterial metabolism under strong selective pressure and highlight the potential of adaptive evolution as a tool for discovering growth-supporting metabolic states that are difficult to predict through rational engineering alone (*22*).

### Central metabolism rewiring

In *E. coli* reorganization of central metabolism following TCA cycle interruption at AKG dehydrogenase appears to be centered on maintaining succinyl-CoA availability (*6*). Consistent with this observation, expression of ACIAD3390, a putative succinyl-CoA acetate CoA-transferase/hydrolase, was strongly increased in ADP1 (Fig. 4) and flux of the reaction increased (Fig. 3C), suggesting increased capacity for succinate/succinyl-CoA interconversion. This supports the hypothesis that succinyl-CoA represents a significant metabolic constraint in TCA-deficient strains due to its essential role in amino acid and cofactor biosynthesis. This interpretation is further supported by intracellular metabolite measurements, which showed strong depletion of succinate, fumarate, and malate despite restoration of growth (Fig. 3A and 3B), indicating persistent depletion of metabolites located downstream of the disrupted AKG-to-succinyl-CoA conversion step. Furthermore, the absence of glyoxylate-shunt flux in the evolved isolates decreased succinate replenishment from isocitrate while the near-zero flux through succinate dehydrogenase limited succinate consumption (Fig. 3C). This outcome was unexpected because glyoxylate-shunt activity is often considered a potential mechanism for conserving carbon and maintaining TCA-cycle intermediates under conditions where oxidative metabolism is constrained. Instead, the evolved isolates adopted a strategy characterized by elimination of glyoxylate-shunt flux together with reduced succinate dehydrogenase activity, suggesting that conservation of succinate-derived intermediates was achieved through redistribution of flux rather than replenishment through the glyoxylate pathway. Similar adaptations have been reported in *E. coli*, where reduced succinate dehydrogenase activity was a major contributor to growth restoration following TCA cycle disruption and in addition reduction of acetyl-CoA entry into the TCA cycle through modulation of *gltA* improved growth but was not essential (*6*). In contrast, restoration of growth in TCA-disrupted ADP1 on glucose minimal medium appears to involve broader reorganization of central metabolism.

The increased growth rate on glucose was accompanied by greater residual carbon at the end of growth (Fig. S3), suggesting that adaptive evolution favored growth rate over complete substrate utilization. Consistent with this shift, metabolic flux analysis revealed reduced carbon entry into the oxidative TCA cycle, increased acetate overflow, and elevated flux through the pyruvate-PEP-OAC-malate node (Fig. 3C). Transcriptomic changes, including upregulation of ED-pathway and acetate-production genes together with decreased expression of acetate assimilation pathways, supported the observed acetate-overflow phenotype (Fig. 4 and Data S4). Notably, acetate secretion has generally not been reported during growth of wild-type ADP1 on glucose, even at elevated substrate concentrations (*23*). Acetate overflow may therefore represent an adaptation to limited oxidative TCA-cycle capacity, enabling disposal of excess acetyl-CoA while conserving energy through substrate-level ATP generation. This interpretation is supported by intracellular metabolite measurements showing accumulation of AKG and glutamate together with depletion of downstream TCA-cycle intermediates (Fig. 3A and 3B), indicating that evolution favored rerouting of carbon around persistent TCA-cycle bottlenecks rather than restoration of balanced TCA-cycle flux. mutations upstream of gltA were accompanied by reduced gltA expression (Table 1 and Fig. 4). Together, these observations suggest reduced carbon flux through citrate synthase despite elevated prpC expression (Data S4), favoring anaplerotic metabolism and acetate overflow. Because increased prpC expression evolved as a compensatory response to reduced citrate synthase activity in an E. coli ΔgltA strain (*21*), elevated prpC expression in the evolved ADP1 isolates may represent a similar adaptation, permitting limited carbon entry into the oxidative branch through the reported secondary citrate synthase activity of PrpC.

In parallel with acetate overflow, the pyruvate-PEP-OAC-malate node appears to play an increasingly important role in carbon redistribution. As a central hub of anaplerotic metabolism, this node replenishes OAC-derived biosynthetic intermediates and may contribute to redox balancing. ^13^C MFA revealed increased flux through this node following TCA-cycle disruption, including elevated conversion of PEP to OAC and reversal of net flux between OAC and malate (Fig. 3C). Increased expression of genes associated with these reactions (Fig. 4) was consistent with the flux analysis and suggests transcriptional reinforcement of this metabolic state. The increased PEP-to-OAC flux, followed by reduction to malate and subsequent conversion to pyruvate, is consistent with enhanced cycling within the PEP-OAC-malate pool, which could sustain precursor supply while facilitating exchange of reducing equivalents. Together, the increased acetate secretion, elevated anaplerotic flux and accumulation of residual carbon support a model in which adaptive evolution favored carbon redistribution through overflow and anaplerotic pathways rather than complete oxidation via the TCA cycle.

### Respiratory and redox adaptation

The reorganization of central metabolism was accompanied by substantial changes in oxidative phosphorylation. The most pronounced differences between the evolved isolates and the reverse-engineered strain were observed in the ETC, particularly in components that transfer electrons to and from the quinone pool. Because both the *nuo* complex and SDH contribute reducing equivalents to the quinone pool, the reduced *nuo* expression (Fig 4) together with lower SDH flux (Fig. 3C) would be expected to decrease electron input to the respiratory chain. The alternative NADH dehydrogenase Ndh, which also transfers electrons from NADH to the quinone pool, was likewise downregulated in the evolved isolates. The downregulation of the *nuo* operon and *ndh* may therefore reflect reduced demand for quinone reduction and respiratory electron transport rather than solely active suppression of respiratory metabolism. This interpretation is consistent with the broader metabolic reorganization observed in the evolved isolates, which reduces carbon oxidation through the TCA cycle while increasing reliance on anaplerotic and overflow pathways. At the same time, increased expression of cytochrome bd oxidase and cytochrome c oxidase may indicate that the terminal step of the ETC is remodeled rather than simply downregulated. Although NADH-dependent quinol production was downregulated, the quinol pool may be sustained by other quinone-reducing pathways, such as sugar oxidation and pyruvate dehydrogenase activity. Together, these changes suggest selective reorganization of quinone-linked electron transport rather than uniform downregulation of aerobic respiration.

A notable feature shared between the metabolic and respiratory adaptations is the strong reduction in succinate dehydrogenase activity. In TCA-deficient *E. coli*, reduced SDH activity contributes to conservation of succinate-derived intermediates and maintenance of succinyl-CoA availability (*6*). However, SDH is also an integral component of the respiratory chain. Adaptive evolution of NADH dehydrogenase *ndh* deficient *E. coli* similarly selected for reduced SDH activity (*24*), indicating that modulation of electron transfer from SDH into the quinone pool can improve respiratory fitness under conditions of altered electron transport. Together with the remodeling of quinone-linked respiratory function expression observed in ADP1, these observations suggest that SDH represents a key interface between central carbon metabolism and aerobic respiration during adaptation to TCA cycle disruption.

The increased flux through the pyruvate-PEP-OAC-malate node in the evolved strains may suggest increased reliance on intracellular redox-balancing reactions (Fig. 3C). This was supported by elevated expression of both malate dehydrogenase genes (*mqo* and *mdh*) (Fig. 4). In addition, conversion of malate to pyruvate through malic enzyme (ACIAD2287) may contribute to NADPH generation, potentially supporting biosynthetic demand. Further evidence for redox remodeling was observed in carbon catabolism, where expression of the NADP-dependent GAP dehydrogenase was strongly increased (Fig. 4), suggesting increased capacity for NADPH generation. A similar pattern was evident at the isocitrate dehydrogenase step. While expression of the canonical isocitrate dehydrogenase *icd* decreased, the alternative isocitrate dehydrogenase *idh* was strongly upregulated (Fig. 4). Because both enzymes catalyze the conversion of isocitrate to α-ketoglutarate with simultaneous reduction of NADP^+^, this reciprocal regulation is consistent with isozyme replacement and suggests altered regulation of NADPH generation rather than complete suppression of the isocitrate-to-AKG conversion step. Together, these observations indicate extensive reorganization of NADPH-producing reactions following TCA cycle disruption. A related shift toward NADPH-generating metabolism has been reported in hydrogen peroxide-stressed *Pseudomonas fluorescens*, where repression of several TCA-cycle enzymes was accompanied by increased activity of NADPH-generating pathways and enhanced pyruvate production (*25*). Although the initiating stress differs from that imposed by TCA-cycle disruption, both systems highlight the importance of redox-oriented metabolic rewiring when oxidative metabolism is constrained.

Changes in NADPH-generating reactions were accompanied by reciprocal regulation of the two transhydrogenase systems linking the NAD(H) and NADP(H) pools. The membrane-bound pyridine nucleotide transhydrogenase *pnt* was strongly downregulated, whereas the soluble transhydrogenase *sthA* was strongly upregulated (Fig. 4, Data S4). Because these enzymes provide alternative routes for exchange of reducing equivalents between NAD(H) and NADP(H), their reciprocal regulation further supports extensive remodeling of cellular redox metabolism. The increased expression of *sthA* may provide greater flexibility in balancing NADH and NADPH demands under conditions where respiratory NADH oxidation is reduced. Simultaneously, downregulation of the PMF-coupled transhydrogenase *pnt* could reduce reliance on PMF-driven NADPH generation, potentially conserving proton motive force under the modified ETC conditions. Together, these observations suggest that enhanced anaplerotic cycling and remodeling of NAD(P)H-producing reactions partially compensate for reduced respiratory electron flux by providing alternative routes for cofactor recycling and redox homeostasis.

Additional evidence for remodeling of quinone-associated electron transfer was provided by physiological responses during fed-batch cultivation. Both ASA557-6 and RE3 resumed growth following glucose or xylose supplementation despite the continued presence of substantial amounts of carbon in the culture medium as downstream oxidation products. Concurrent decreases in dissolved oxygen further indicated renewed respiratory activity during periods of resumed growth (Fig. S4). These observations suggest that carbon availability alone was not limiting growth. Rather, continued biomass formation may have depended on renewed electron delivery to the respiratory chain under TCA-constrained conditions. The ability of both strains to also resume growth following formate supplementation further supports the importance of electron-donor availability, as formate oxidation can provide reducing equivalents while contributing little carbon to biomass formation. Consistent with this interpretation, expression of the pyrroloquinoline quinone (PQQ)-dependent glucose dehydrogenase (*gcd*), which transfers electrons from periplasmic glucose oxidation into the respiratory ETC via the quinone pool, was modestly increased in the evolved isolates but remained near reference levels in the reverse-engineered strain (Fig. 4), suggesting enhanced utilization of quinone-linked electron-input pathways in the evolved backgrounds. Although the evolved isolates and RE3 exhibited similar growth rates, the evolved isolates consistently achieved higher final biomass concentrations (Fig. 2B and Fig. S3). The more extensive remodeling of quinone-linked electron transport in the evolved strains therefore correlates with improved biomass formation under TCA-constrained conditions, consistent with an enhanced capacity to sustain respiratory electron flux, although the precise mechanisms underlying this advantage remain unclear.

### Biosynthetic reallocation

Although growth was restored, intracellular pools of several amino acids remained depleted relative to the reference strain (Fig. 3A and 3B). In particular, aspartate and glycine remained at low levels despite transcriptional upregulation of biosynthetic pathways originating from oxaloacetate and 3-phosphoglycerate in ASA557-6, suggesting that precursor demand continued to exceed supply under TCA-constrained conditions. Elevated flux through the pyruvate-PEP-OAC-malate node likely alleviated, but did not fully overcome, these limitations. Similar decoupling between metabolic flux, growth-associated functions, and biosynthetic regulation has recently been observed in bacteria adapting to physiological perturbations (*26*). Accumulation of AKG and glutamate was likely driven by disruption of the α-ketoglutarate dehydrogenase step, which blocks conversion of AKG to succinyl-CoA and alters utilization of the surrounding metabolite pool (Fig. 3A and 3B). Notably, elevated AKG and glutamate levels persisted despite restoration of growth, indicating that adaptive evolution enabled growth within a substantially altered metabolic landscape rather than restoring pre-perturbation metabolite pools. While expression of most pathways consuming these intermediates remained low, upregulation of arginine and cyanophycin metabolism (Data S4) may have contributed to redistribution of glutamate-derived carbon and nitrogen. In contrast, repression of branched-chain amino acid biosynthesis, together with low intracellular levels of leucine and isoleucine, may reflect reduced investment in biosynthetically expensive pathways, thereby easing demands on carbon and reducing-equivalent availability.

Genes involved in fatty acid biosynthesis were among the most strongly upregulated pathways (Fig. 4), consistent with increased demand for membrane lipid synthesis during rapid growth (Fig. S3). Increased expression of genes responsible for malonyl-CoA formation and fatty acid elongation, together with reduced expression of *fadD*, suggests net allocation of carbon toward lipid biosynthesis rather than fatty acid turnover. Because fatty acid synthesis is a major sink for both acetyl-CoA and NADPH, its upregulation implies increased demand for carbon precursors and reducing power. Concurrent upregulation of biotin biosynthesis genes further supports increased acetyl-CoA carboxylase activity and malonyl-CoA production. Together, these changes are consistent with a shift toward growth-associated biosynthesis despite reduced carbon-use efficiency. Notably, depletion of multiple TCA-derived intermediates and amino acids alongside elevated AKG and glutamate levels (Fig. 3A and 3B) indicates that improved growth was not accompanied by restoration of metabolic homeostasis. Instead, adaptation appears to have maintained biosynthetic capacity and metabolic throughput despite persistent perturbations in central metabolism.

### Regulatory adaptation and reverse engineering

In contrast to *E. coli*, where adaptation to severe TCA-cycle disruption can largely be achieved through modifications affecting TCA-cycle flux and succinate metabolism (*6*), equivalent changes were insufficient to restore growth in ADP1 on glucose minimal medium (Fig. 2B). These differences suggest that restoration of growth in a strictly aerobic bacterium requires broader reorganization of metabolism and respiratory physiology than in facultative organisms. The extensive rewiring observed in central metabolism, oxidative phosphorylation, redox balancing and biosynthetic allocation indicates that adaptation extends beyond correction of a single metabolic bottleneck.

Mutations in both the two-component regulatory system *qseBC* and *ACIAD2336* were required for restoration of growth in the reverse-engineered strain (Fig. 2B), highlighting their importance for adaptation. *ACIAD2336* encodes a putative ATPase homologous to the cell division factor ZapE of *E. coli* (*27*). Although the physiological consequences of the mutation remain unclear, ZapE-family proteins are associated with stress-responsive cell division and maintenance of cytokinesis under challenging growth conditions. Alteration of ACIAD2336 may therefore have facilitated growth under the substantial metabolic and physiological perturbations imposed by TCA-cycle disruption. Similarly, the nonsense mutation identified in *qseC,* which was also accompanied by reduced *qseB* expression (Data S4), indicating substantial perturbation of the QseBC regulatory circuit, may have contributed to adaptation through broader effects on cellular physiology. Beyond its established roles in environmental sensing and quorum-regulated behaviors, QseBC functions as a global regulator of bacterial physiology. Studies in *E. coli* and *Aggregatibacter actinomycetemcomitans* have shown that QseBC influences central metabolism, respiration, energy production, and adaptation to environmental conditions, with *qseC* disruption in *E. coli* causing broad perturbations in amino acid metabolism and the TCA cycle (*28*, *29*). These observations suggest that disruption of QseBC in ADP1 may have facilitated adaptation by relaxing native regulatory programs controlling metabolism and growth, thereby enabling establishment of an alternative metabolic state following TCA-cycle disruption. Together, these findings suggest that restoration of growth required changes not only in metabolism but also in the cellular systems coordinating growth and stress adaptation.

Despite restoring growth on glucose minimal medium, the reverse-engineered strain did not fully reproduce the phenotype of the evolved isolates, particularly with respect to final biomass accumulation (Fig. 2B and Fig. S3). This observation indicates that additional mutations and epistatic interactions contribute to the adapted state. Several evolved mutations were identified in genes associated with transcriptional and translational processes, including *rpoC*, *greA*, and *rpsF* (Table 1). Although these genes did not exhibit pronounced transcriptional responses, their effects are likely manifested through altered transcriptional regulation, translation efficiency, or other post-transcriptional mechanisms. In particular, mutations affecting RNA polymerase subunits are frequently selected during adaptive laboratory evolution and can facilitate adaptation through global transcriptional reprogramming rather than pathway-specific regulation. Previous studies demonstrated that *rpoC* mutations improved growth in minimal media by altering RNA polymerase kinetics, redistributing transcriptional resources, and reshaping genome-wide gene expression patterns (*30*). The *rpoC* mutation identified here may therefore have contributed to adaptation by adjusting transcriptional priorities to better support the rewired metabolic and respiratory state of the evolved strains. Similarly, GreA functions as a transcription elongation factor that promotes recovery of stalled RNA polymerase complexes and reduces promoter-proximal pausing (*31*, *32*). Mutations affecting *greA* may therefore influence transcriptional dynamics across large portions of the genome and facilitate adaptation to the extensive physiological perturbations imposed by TCA-cycle disruption. The mutation identified in *rpsF*, which encodes the ribosomal protein S6, may likewise reflect adaptation at the level of translational resource allocation. Because RpsF is a component of the translation machinery and participates in the regulation of ribosome biogenesis, alterations affecting its function could improve coordination between protein synthesis capacity and the altered metabolic demands of the evolved strains (*33*). In contrast, the histone-like protein HupB was consistently downregulated, suggesting that adaptation was also accompanied by changes in chromosome organization and global gene regulation. Collectively, these results indicate that adaptation to TCA-cycle disruption emerged through multilayered regulatory evolution, in which metabolic rewiring was accompanied by broader changes in cellular physiology and gene regulation.

### Selection constraints, adaptive states, and outlook

Successful adaptation required careful selection of evolutionary pressure. Restoration of growth was only achieved under specific ALE conditions despite testing multiple cultivation strategies (Fig. S1), indicating that accessibility of adaptive metabolic states was strongly influenced by the selection environment. Conditions designed to promote glyoxylate-shunt activity, such as acetate supplementation, did not yield the expected phenotype and instead opposed the metabolic state that ultimately emerged, which was characterized by acetate overflow and reduced reliance on the glyoxylate pathway (Fig. 3C). Likewise, succinate supplementation alleviated the selective pressure imposed by TCA-cycle disruption and reduced the need for the extensive flux redistribution observed around the succinate-succinyl-CoA node. In contrast, gradual reduction of yeast extract maintained growth while progressively increasing dependence on endogenous glucose metabolism, enabling the adaptive trajectory observed in the evolved isolates. These observations highlight the importance of balancing growth support and selective pressure when designing adaptive evolution experiments (*34*).

Consistent with the multilayered regulatory and metabolic adaptations discussed above, the reverse-engineered strain did not fully reproduce the evolved phenotype. Although reconstruction of key mutations restored growth on glucose minimal medium, additional mutations and epistatic interactions contributed to the final phenotype, particularly with respect to biomass accumulation. These results demonstrate that adaptation to severe TCA-cycle disruption extends beyond correction of individual metabolic bottlenecks and instead involves coordinated reorganization of metabolism, respiration, redox balancing, biosynthesis, and cellular regulation. Compared with previous studies on restoration of growth following TCA-cycle disruption the strictly aerobic bacterium ADP1 required substantially broader physiological reprogramming than *E. coli* (*6*). In particular, remodeling of quinone-linked electron transport emerged as a prominent feature of the adapted phenotype, distinguishing the evolved isolates from the reverse-engineered strain and highlighting the importance of respiratory adaptation following disruption of central carbon metabolism. However, the mechanistic basis of several key adaptations remains unresolved. Future work should therefore focus on determining the contribution of additional mutations and their interactions with the reconstructed genotype. Direct measurements of respiratory activity, oxygen uptake, intracellular cofactor ratios, and quinone-pool redox state will be required to fully resolve how metabolic, respiratory, and regulatory changes collectively support growth under TCA-constrained conditions.

From a metabolic engineering perspective, disruption of the TCA cycle effectively redirected carbon away from oxidative metabolism and toward biosynthetic and anaplerotic pathways. However, this redistribution was accompanied by substantial acetate overflow, demonstrating that carbon not oxidized through the TCA cycle can instead be lost through overflow metabolism. Consequently, TCA-cycle engineering alone may not be sufficient to maximize carbon retention in aerobic production hosts. Combining TCA-cycle disruption with strategies that limit acetate formation or promote acetate reassimilation may therefore provide a more effective route toward improving carbon utilization. Overall, the simplified architecture of glucose metabolism in ADP1 facilitated identification of the physiological, metabolic and regulartory adaptations required to restore growth following severe TCA-cycle perturbation, highlighting its value as a model chassis for studying large-scale metabolic rewiring. More broadly, these results demonstrate that adaptive laboratory evolution can uncover growth-supporting metabolic states that are difficult to predict through rational engineering alone and reveal alternative growth-supporting metabolic configurations that become accessible following major perturbations to central metabolism.

## MATERIALS AND METHODS

### Strains and Media

*A. baylyi* ADP1 (DSM 24193, DSMZ, Germany) was used as the host for all the experiments. All the strains used in the study are listed in Table 2.

**Table 2.** Strains used in this study.

| Strain name | Organism | Genotype or description | Reference of source |
| --- | --- | --- | --- |
| ASA549 | *A. baylyi* ADP1 | $\Delta lldP-dld::ara$ gene cluster I, $\Delta acoA-budC::ara$ gene cluster II | (16) |
| ASA555 | *A. baylyi* ADP1 | $\Delta lldP-dld::ara$ gene cluster I, $\Delta acoA-budC::ara$ gene cluster II, $\Delta sucAB$, $\Delta gabT::tdk-kan^r$ | (16) |
| ASA557 | *A. baylyi* ADP1 | Single colony isolated from the evolution lines. Six individual colonies, designated as ASA557-1 to ASA557-6 | This study |
| Rescued ASA555 | *A. baylyi* ADP1 | $\Delta lldP-dld::ara$ gene cluster I, $\Delta acoA-budC::ara$ gene cluster II, $\Delta sucAB$, $\Delta gabT$ | This study |
| ASA558 (RE1) | *A. baylyi* ADP1 | Reverse-engineered strain. The rescued ASA555 with deletion of the *sdhA-gltA* region and replacement by region carrying the *sdhA* (S300T) and *gltA* upstream mutations at the native locus | This study |
| ASA559 (RE2) | *A. baylyi* ADP1 | Reverse-engineered strain. ASA558 with *qseC* replaced by *qseC* (Q219*) | This study |
| ASA560 (RE3) | *A. baylyi* ADP1 | Reverse-engineered strain. ASA559 with the deletion of the region spanning *glcB* to *ACIAD2336* | This study |

Mineral salts medium (MSM) was used for genetic modification and all physiological experiments. Additional components, such as carbon sources (glucose, yeast extract, sodium succinate, sodium acetate, etc.) and the buffering agent MOPS, were added as specified in the Results and other sections. The composition of MSM is 3.88 g/L K_2_HPO_4_, 1.63 g/L NaH_2_PO_4_, 2.00 g/L (NH_4_)_2_SO_4_, 0.1 g/L MgCl_2_·6H_2_O, 10 mg/L ethylenediaminetetraacetic acid (EDTA), 2 mg/L ZnSO_4_·7H2O, 1 mg/L CaCl_2_·2H_2_O, 5 mg/L FeSO_4_·7H_2_O, 0.2 mg/L Na_2_MoO_4_·2H_2_O, 0.2 mg/L CuSO_4_·5H_2_O, 0.4 mg/L CoCl_2_·6H_2_O, and 1 mg/L MnCl_2_·2H_2_O. Kanamycin was added at 50 μg/mL when required.

### Adaptive Laboratory Evolution

Adaptive laboratory evolution (ALE) was applied to restore growth of the TCA-cycle-deficient strain ASA555 on glucose. ALE was carried out in 100-mL Erlenmeyer flasks containing 10 mL of MSM supplemented with glucose and yeast extract, and 50 μg/mL kanamycin was included to prevent contamination. Cultures were incubated at 30°C and 300 rpm. Cells were transferred daily to fresh medium. The OD600 was measured before each transfer, and the inoculum volume was adjusted to give an initial OD of 0.01–0.05. Yeast extract and glucose were initially supplied at 2% (w/v) and 0.4% (w/v), respectively. Over the course of the evolution, the yeast extract concentration was decreased while the glucose concentration was increased, as detailed in the Results section. For the evolution line in which the population gained the ability to grow on glucose as the sole carbon source, yeast extract was eventually omitted, leaving glucose as the sole carbon source. Other ALE strategies tested included supplementation with yeast extract (2% w/v) + glucose (1% w/v) + sodium acetate (40 mM) and sodium succinate (5–25 mM) + glucose (0.4% w/v).

The number of generations (n) per flask was calculated with the following equation:

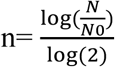

where N is the final OD600 of the culture and N0 is the initial OD600.

After evolution, single clones were isolated on MSM agar plates containing 4 g/L glucose.

### Mutation Analysis

To identify mutations that arose during ALE, whole-genome sequencing was performed on the parental strain ASA555 and the evolved isolates ASA557-1 to ASA557-6. Genomic DNA was extracted using the GeneJET Genomic DNA Purification Kit (Thermo Fisher Scientific). DNA libraries were prepared with the Native Barcoding Kit 24 V14 (Oxford Nanopore Technologies, UK) according to the manufacturer’s instructions. Sequencing was performed on a MinION Mk1B device (Oxford Nanopore Technologies, UK). Reads from each strain were mapped to the ASA555 reference genome (Data S5) derived from GenBank entry CR543861. Mutation calling was performed using breseq 0.38.3 with the mode for nanopore sequencing data (*35*). Large structural rearrangements, such as inversions, were validated by de novo assembly using a custom script. This assembly workflow consisted of sequential Flye assembly, Minimap2-Racon polishing, Medaka consensus refinement, and Prokka annotation.

### Genetic Engineering

To validate the role of key mutations, specific mutations were introduced sequentially into the parental strain ASA555. All primers used in this study are listed in Data S1. Transformation and homologous recombination-based genome editing of ADP1, as well as counter-selection for marker removal, were performed as described previously (*36*). Genomic modifications were made using linear cassettes carrying homologous flanking sequences (1–1.2 kb) generated by overlap extension PCR.

To construct ASA558 (RE1), the kanamycin resistance marker in ASA555 was first removed via counter-selection using a cassette generated from two fragments amplified with primer pairs P1/P2 and P3/P4. Transformation and selection at this step were performed on MSM supplemented with 30 mM succinate. The marker-free strain was then subjected to deletion of the *sdhA*-*gltA* region using a cassette assembled from three fragments: a left flank (amplified with primer pair P5/P6), the *tdk*/*kan*^r^ cassette (P7/P8), and a right flank (P9/P10). For this and all subsequent steps, transformation and selection were performed on MSM supplemented with 1% (w/v) yeast extract and 0.6% (w/v) glucose. The resulting strain was then “rescued” by replacing the deletion cassette with the *sdhA*-*gltA* region amplified from the genome of evolved isolate ASA557-1 using primers P5/P10; this region carries both the *sdhA* (S300T) and *gltA* upstream mutations, yielding strain ASA558 (RE1).

To construct ASA559 (RE2), the *qseC* gene was first deleted in ASA558 (RE1) using a cassette assembled from a left flank (P11/P12), the *tdk*/*kan*^r^ cassette (P13/P14), and a right flank (P9/P10). The deletion cassette was subsequently replaced with the *qseC* region amplified from ASA557-1 using primers P11/P14, introducing the Q219* mutation and generating strain ASA559 (RE2).

To construct ASA560 (RE3), the region spanning *glcB* to *ACIAD2336* was deleted in ASA559 (RE2) using a cassette assembled from a left flank (P15/P16), the tdk/kan^r^ cassette (P7/P8), and a right flank (P17/P18), yielding strain ASA560 (RE3).

### Growth Characterization

Growth of evolved isolates and reverse-engineered strains was characterized in 96-well plates. Cells were cultured in 200 μL of medium in flat-bottom 96-well plates (μClear^TM^, white, Greiner) at 30°C in a Spark multimode microplate reader (Tecan, Switzerland). Double orbital shaking (6 mm amplitude, 54 rpm) was applied for 5 min twice per hour, and OD at 600 nm was measured hourly. For growth profiling of evolved isolates, MSM was supplemented with 0.27% (w/v) glucose (15 mM). For reverse-engineered strains, MSM was supplemented with 0.6% (w/v) glucose (33.3 mM).

For comparison of the evolved isolates ASA557-3 and ASA557-6 with the reference strain ASA549, cells were cultivated in 100-mL Erlenmeyer flasks containing 10 mL MSM with 10 mM (1.8 g/L) glucose as the sole carbon source and 50 mM MOPS as a buffering agent, at 30°C and 300 rpm.

Bioreactor cultivation of ASA557-3 was performed in a mini bioreactor system (Applikon MiniBio). Cells were pre-cultured in MSM supplemented with 1% (w/v) glucose and then inoculated into the bioreactor containing 50 mL MSM with 0.8% (w/v) glucose. Xylose was fed at the following time points and concentrations: 3.3 mM (0.5 g/L) at 17 h, 18.5 h, and 22.5 h; 13.3 mM (2 g/L) at 24.5 h; and 8.3 mM (1.25 g/L) at 38.5 h. Biomass was monitored by OD600 measurement and real-time backscatter using a Cell Growth Quantifier (CGQ). The stirring speed was constant at 200 rpm, and pH was maintained at 6.5 with NaOH.

### Analysis of Substrates and Extracellular Metabolites

Substrate and extracellular metabolite concentrations were determined by high-performance liquid chromatography (HPLC). Acetate, α-ketoglutarate, D-glucose, D-xylose, and L-arabinose were analyzed on an LC-20AC prominence liquid chromatograph (Shimadzu) equipped with an RID-10A refractive index detector. Separation was performed on a Phenomenex Rezex RHM-monosaccharide H⁺ (8%) column with 5 mM sulfuric acid as the mobile phase at a flow rate of 0.5 mL/min and 40 °C.

When sulfuric acid was used, peaks of sugars and sugar acids overlapped. Therefore, in samples containing both, D-glucose and D-xylose were quantified using ultrapure water as the mobile phase. Sugar acid concentrations were estimated as follows: sugar concentrations were first determined from the ultrapure water chromatography. The corresponding peak areas for sugars when using 5 mM sulfuric acid were then predicted from a standard curve, and the sugar acid peak areas were calculated by subtracting the estimated sugar peak areas from the total peak areas obtained under sulfuric acid elution.

### ^13^C-metabolic Flux Analysis

For ^13^C metabolic flux analysis, parallel labeling experiments were performed with [1-^13^C]D-glucose and a 1:1 mixture of [U-^13^C]D-glucose and natural glucose, using strains ASA549 and ASA557-6. Cells were pre-cultured in MSM supplemented with 2 g/L glucose, harvested by centrifugation (3,000 rpm, 5 min), washed with MSM, and resuspended in 10 mL MSM containing the labeled substrates (2 g/L) and 50 mM MOPS in 100-mL flasks. To minimize the contribution of unlabeled biomass, the initial OD was adjusted to 0.01. Cultivation was performed at 30°C and 300 rpm. Samples were periodically withdrawn for OD measurement and HPLC analysis of substrates and extracellular metabolites. When the OD reached 0.45, 0.5 mL samples were collected for analysis of proteinogenic amino acid labeling.

Proteinogenic amino acids were analyzed following a published protocol (*37*). Briefly, cell pellets were hydrolyzed in 500 μL of 6 N HCl at 110 °C for 12–18 h. After hydrolysis, debris was removed by centrifugation, and 400 μl of the supernatant was transferred to a new tube and evaporated at 65 °C under a stream of air or nitrogen. The dried residue was derivatized with 35 μL of pyridine and 50 μL of MTBSTFA + 1% (wt/wt) TBDMSCl (Sigma-Aldrich, cat. no. 375934) at 60 °C for 30 min, and 75 μL of the derivatized sample was transferred to a vial for GC-MS analysis. GC-MS was performed on a GCMS-QP2020 NX gas chromatograph-mass spectrometer (Shimadzu) equipped with a Nexis GC-2030 system and a Zebron ZB-5MSi column (30 mm × 0.25 mm × 0.25 μm, Phenomenex). The injection volume was 1 μL, with helium as carrier gas (1 mL/min). The temperature program was: 80 °C (2 min), ramp to 280 °C at 7°C/min, hold 20 min. Isotopomer distributions were corrected for natural isotope abundance using IsoCor (*38*). Corrected data are provided in Data S3. Flux estimation was carried out with influx_s (*39*), using a central metabolic model constrained by the mass isotopomer distributions, specific growth and substrate uptake rates, and extracellular product secretion rates. The metabolic network of ADP1 constructed previously (*16*) was used. Confidence intervals for fluxes were calculated by influx_s based on Monte Carlo simulations. Complete MFA results are available in Data S2.

### Quantification of Intracellular Metabolites

To quantify intracellular metabolites in ASA549 and ASA557-6, cells were cultivated in MSM with 2 g/L glucose and 50 mM MOPS. Culture samples (10 mL) were taken at OD600 of approximately 0.55 and 0.95, centrifuged at 3000 rpm at 4°C, and the pellets were immediately extracted using a protocol adapted from (*37*). Briefly, 1 mL of preheated 70% (vol/vol) ethanol was added to the pellet and vortexed for 30 s, followed by incubation at 95 °C for 5 min. After centrifugation, the supernatant was collected and evaporated to dryness at 37 °C. The dried residue was derivatized by adding 50 μL of 2% (wt/vol) methoxylamine hydrochloride in pyridine and incubating at 37 °C for 90 min, followed by addition of 50 μL of MTBSTFA + 1% (wt/wt) TBDMSCl (Sigma-Aldrich, cat. no. 375934). After centrifugation, 75 μL of the supernatant was transferred to a GC-MS vial. The GC program was the same as for ^13^C-metabolic Flux Analysis.

A standard mixture containing alanine, glycine, leucine, isoleucine, succinate, proline, fumarate, methionine, serine, α-ketoglutarate, threonine, phenylalanine, malate, aspartate, glutamate, citrate, and tyrosine was prepared at concentrations listed in Table S1. The standard was dried under airflow and subjected to the same procedure as the samples. Calibration curves as specified in Table S1 were used to calculate the abundance of each metabolite.

### Transcriptomics Analysis

For whole transcriptome analysis strains ASA549, ASA557-3, ASA557-6 and RE3 were cultivated in MSM with 2 g/L glucose and 50 mM MOPS. Samples were taken at OD600 of approximately 0.35 and centrifuged at 5000 x g 4°C and washed once with 1X PBS and the final cell pellet was stored at −80 °C. RNA extraction and paired-end Illumina sequencing was performed by Novogene Co., Ltd. using the Illumina NovaSeq platform with 150 bp paired-end reads. The raw sequencing data quality control was done by running fastQC 0.12.1 before and after trimming with fastp 1.3.1 (*40*). Trimmed reads were aligned to the ASA549 genome derived from GenBank entry CR543861 using Bowtie2 2.5.5 (*41*) and gene level read counts were generated with featureCounts 2.1.1 (*42*).

### Statistical Analysis

Differences in metabolite abundance between ASA549 and ASA557-6 were assessed using Student’s t-test (two-tailed, three replicates per group). Significance levels are indicated as * p < 0.05, **p < 0.01, ***p < 0.001.

For metabolic flux analysis, model fitting was evaluated using the χ^2^ (chi-square) goodness-of-fit test. After fitting, the χ^2^ statistic and its corresponding *p*-value were calculated. A *p*-value ≥ 0.05 indicated that the null hypothesis, namely that the model adequately describes the experimental data, could not be rejected; therefore, the fit was considered statistically acceptable.

Confidence intervals for estimated fluxes were determined using a Monte Carlo approach. Synthetic measurement datasets were generated by adding random noise to the original experimental measurements, and fluxes were re-estimated for each dataset. The 2.5^th^ and 97.5^th^ percentiles of the resulting flux distributions were then used to define the 95% confidence intervals.

Differential expression analysis was performed with DESeq2 1.52.0 (*43*). Raw gene counts were normalized using DESeq2’s median-of-ratios method to account for differences in sequencing depth and RNA composition among samples. Differential expression was assessed by fitting a negative binomial generalized linear model for each gene and testing model coefficients using Wald tests. Resulting P-values were corrected for multiple hypothesis testing using the Benjamini-Hochberg false discovery rate (FDR) procedure. To improve effect-size estimation, log2 fold-change values were shrunken using apeglm 1.34.0 (*44*) using DESeq2’s lfcShrink() function. Genes were called differentially expressed at adjusted P-value < 0.05 and absolute shrunken log2 fold-change ≥ 0.5. Variance-stabilizing transformation implemented in DESeq2 was used for exploratory analyses and visualization of sample relationships and principal component analysis was performed on variance-stabilized count data.

## Author contributions

Conceptualization: JL, ET, CL

Methodology: ET, CL, JL

Investigation: CL, ET, JL

Visualization: ET, JL

Supervision: VS, SS, CL, JL

Writing—original draft: CL, ET, JL Writing—review & editing: ET, CL, JL, VS, SS

## Data and materials availability

The data supporting the findings of this study are in the main text or the supplementary materials. The corresponding author is willing to provide the raw data related to this manuscript upon reasonable request. FASTQ files from genome sequencing of ASA555 and ASA557-1 to ASA557-6 are available from the NCBI Sequence Read Archive (PRJNA1530000).

Transcriptomics analysis code is available in the Github repository: https://github.com/Eemeli-T/RNA-sequencing-bioinformatics

