## Supplementary materials for "Metabolic adaptation to TCA cycle deficiency in a strictly aerobic bacterium"

Changshuo Liu *et al.*

**This PDF file includes:**

Figs. S1 to S10
Table S1

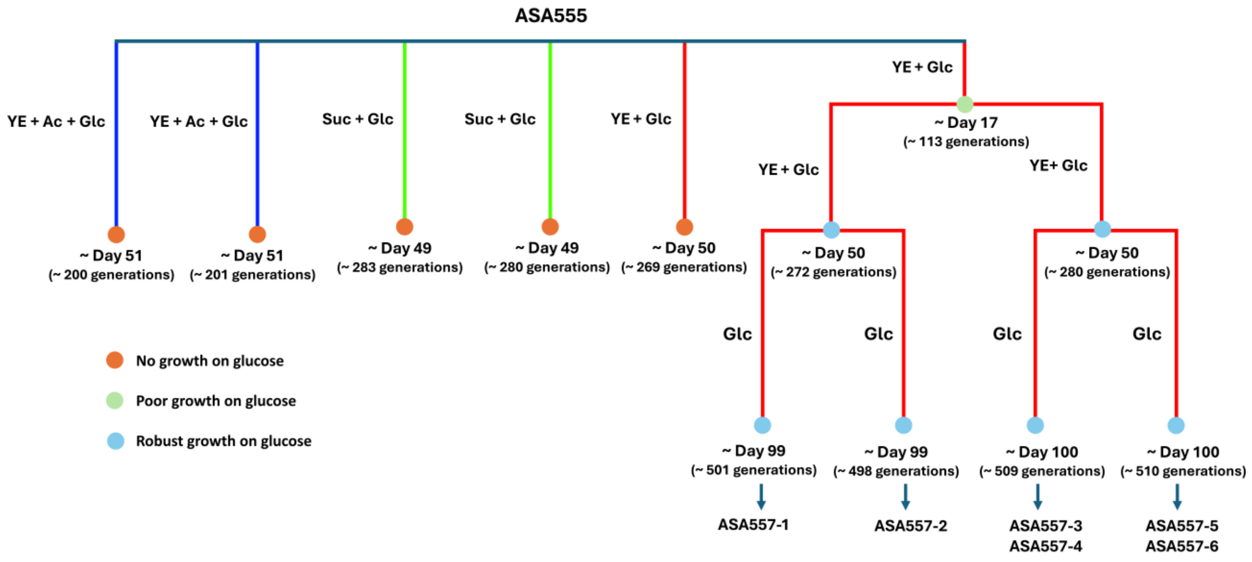


**Fig. S1.**

Growth outcomes (indicated with colored circles) and evolution trajectories of ASA555 under different adaptive evolution conditions. Cells were adapted in mineral salts medium supplemented with various carbon source combinations: yeast extract plus acetate and glucose (YE + Ac + Glc), succinate plus glucose (Suc + Glc), yeast extract plus glucose (YE + Glc), or glucose alone (Glc). Evolution lines belonging to the same experiment group are indicated in the same color. Abbreviations: YE, yeast extract; Ac, acetate; Glc, glucose; Suc, succinate.

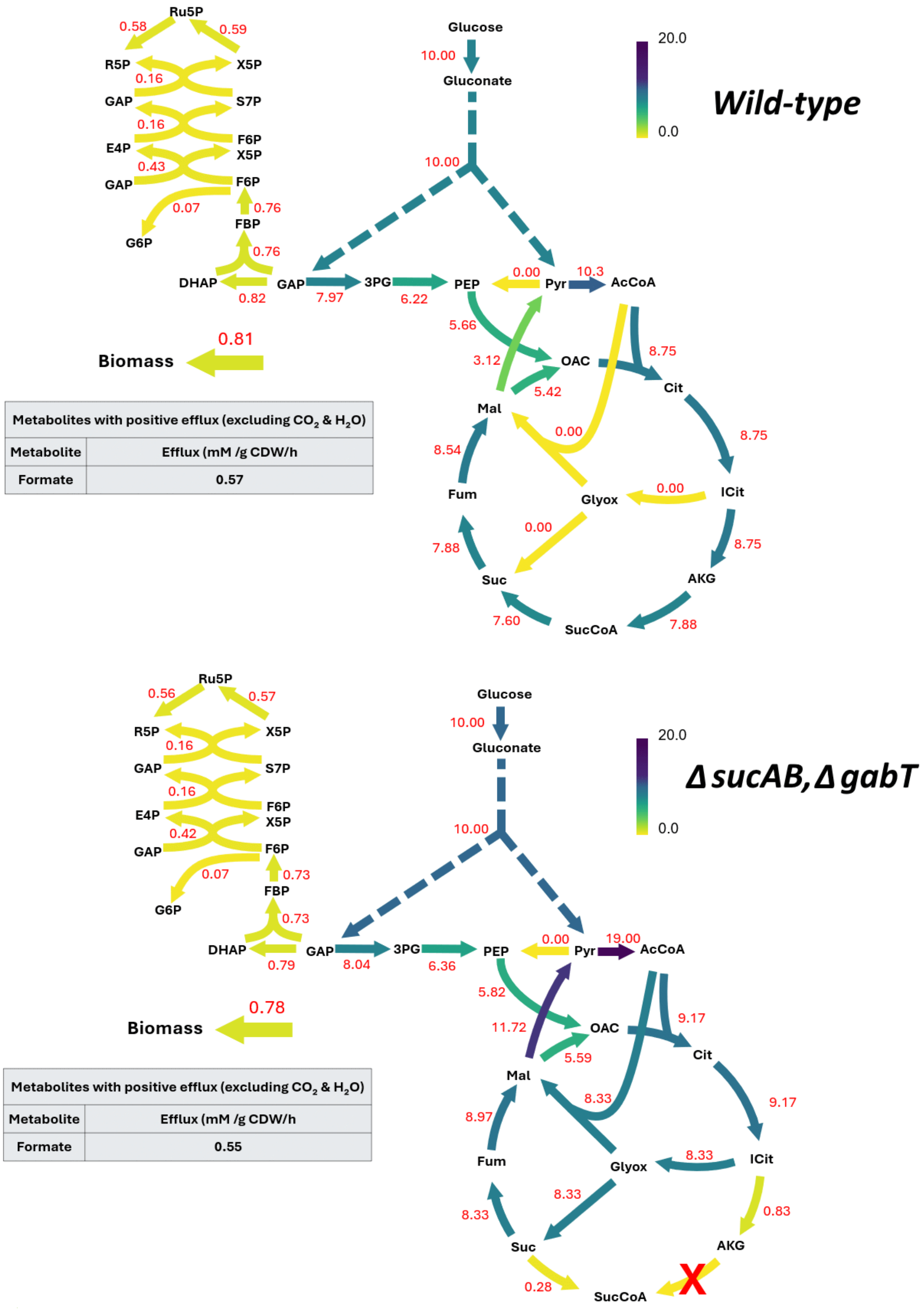


**Fig. S2.**

Flux distributions predicted by flux balance analysis (FBA) for the wild-type and TCA cycle-deficient strain (Δ*sucAB*, Δ*gabT*). Glucose uptake rate was set to 10 mmol/gCDW/h, with all other nutrients assumed to be in excess. The biomass production reaction was used as the objective function.

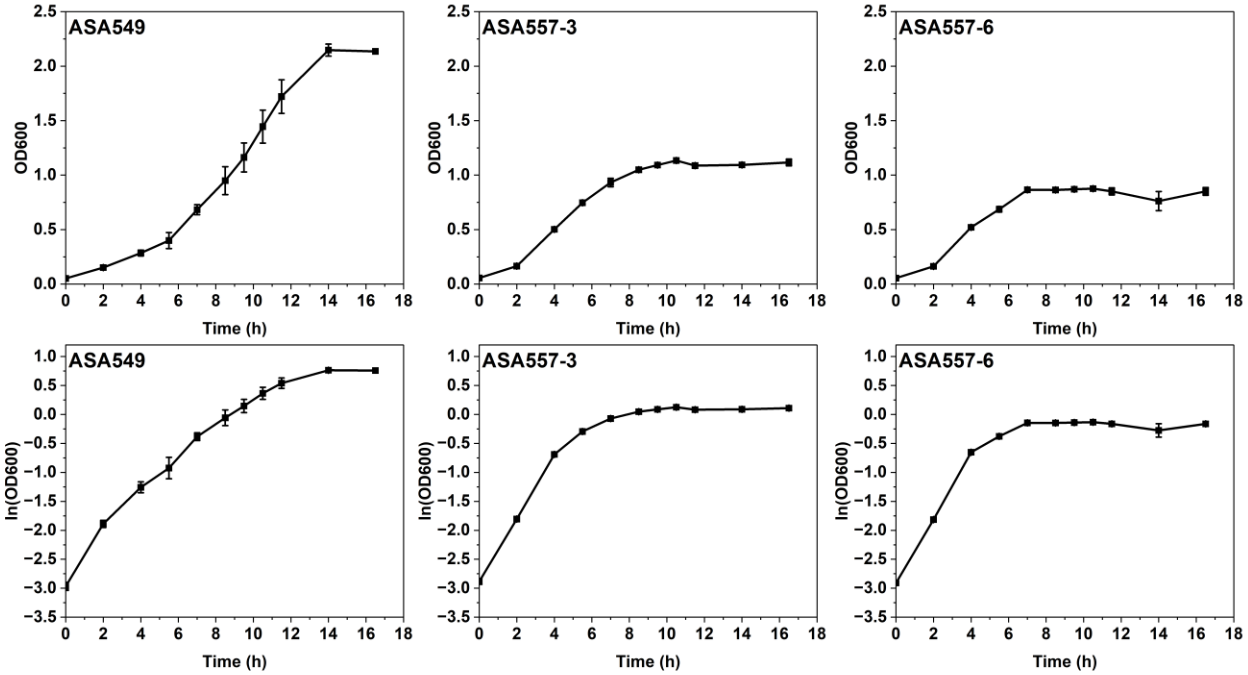


**Fig. S3.**

Growth of strain ASA549 and evolved isolates ASA557-3 and ASA557-6 on glucose. Cells were cultivated in mineral salts medium supplemented with 10 mM (1.8 g/L) glucose as the sole carbon source. Data represent mean values and error bars indicate standard deviations from two independent biological experiments.

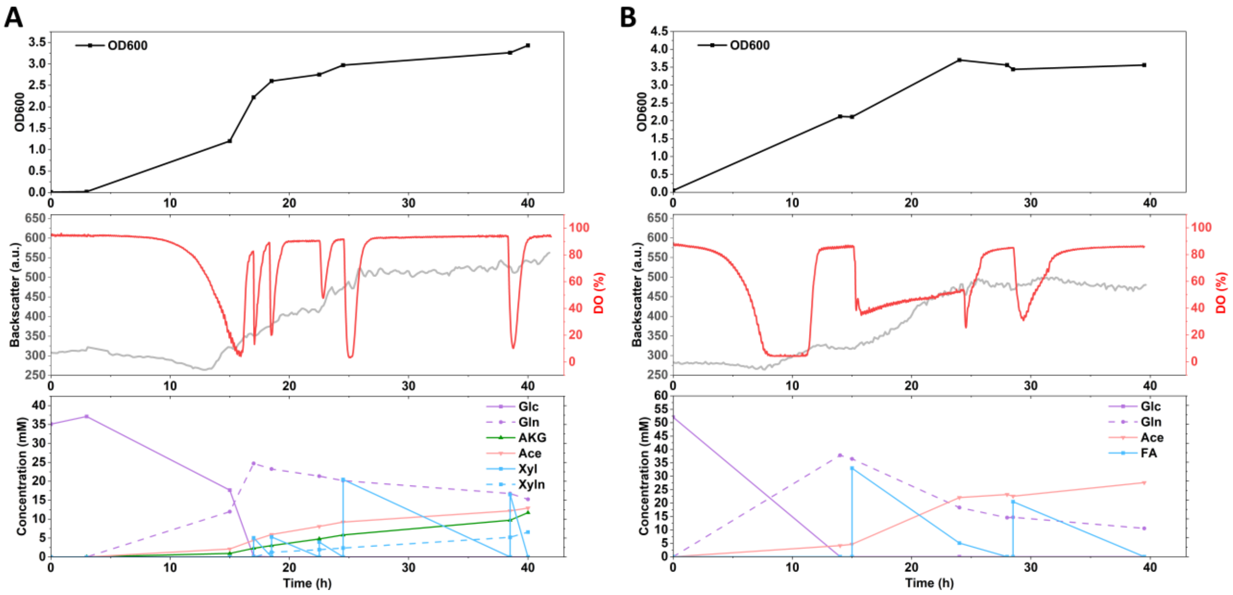


**Fig. S4.**

Bioreactor cultivation of the evolved isolates ASA556-3 and ASA556-6. (**A**) Fed-batch cultivation of ASA556-3 with xylose supplementation. Cells were grown in mineral salts medium containing glucose. Xylose was fed at the following time points and concentrations: 3.3 mM (0.5 g/L) at 17 h, 18.5 h, and 22.5 h; 13.3 mM (2 g/L) at 24.5 h; and 8.3 mM (1.25 g/L) at 38.5 h. (**B**) Fed-batch cultivation of ASA556-6 with formic acid supplementation. Cells were grown in mineral salts medium containing glucose. Formic acid was fed at the following time points and concentrations: 20 mM at 15 h; 10 mM at 28.5 h. The pH was maintained at 7.0, and the stirring speed was fixed at 200 rpm. Optical density at 600 nm (OD600), dissolved oxygen (DO), and extracellular metabolite concentrations were monitored. Backscatter was measured in real time using a Cell Growth Quantifier (CGQ) and correlated with optical density. Abbreviations: Glc, glucose; Gln, gluconate; AKG, α-ketoglutarate; Ac, acetate; Xyl, xylose; Xyln, xylonate; FA, formic acid.

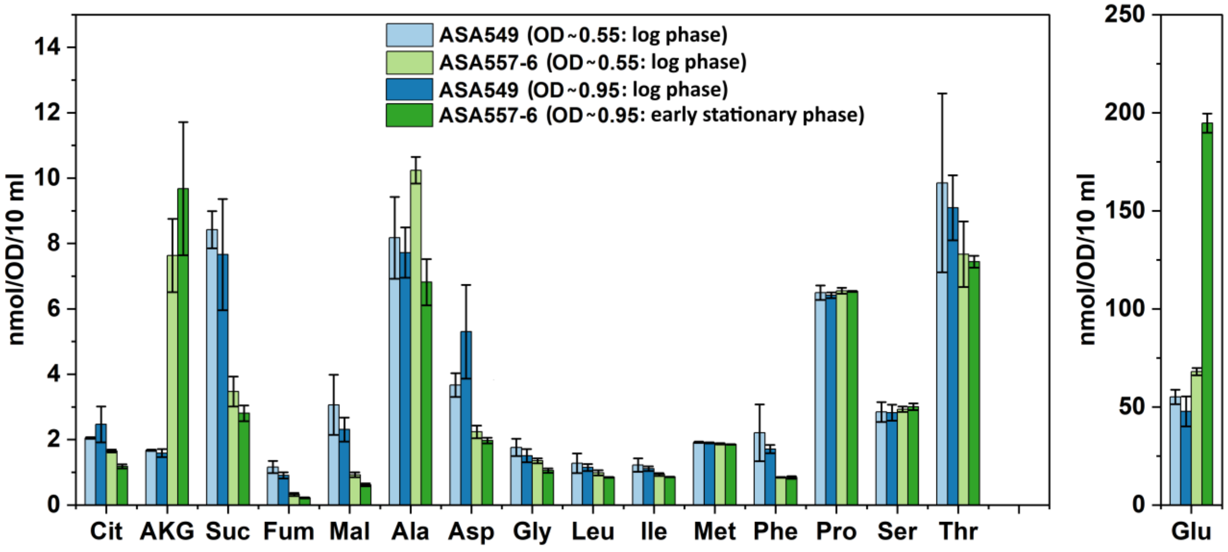


**Fig. S5.**

Quantification of intracellular metabolites in strains ASA549 and ASA557-6 by GC-MS. Cells were cultivated in mineral salts medium supplemented with 11.1 mM (2 g/L) glucose and 50 mM MOPS. Equivalent biomass samples were harvested for metabolite extraction at two time points: (1) OD600 ≈ 0.55 (log phase for both strains) and (2) OD600 ≈ 0.95 (log phase for ASA549; early stationary phase for ASA557-6). Data represent mean values and error bars indicate standard deviations from three independent biological experiments. Abbreviations: Cit, citrate; AKG, α-ketoglutarate; Suc, succinate; Fum, fumarate; Mal, malate; Ala, alanine; Asp, aspartate; Gly, glycine; Leu, leucine; Ile, isoleucine; Met, methionine; Phe, phenylalanine; Pro, proline; Ser, serine; Thr, threonine; Glu, glutamate.

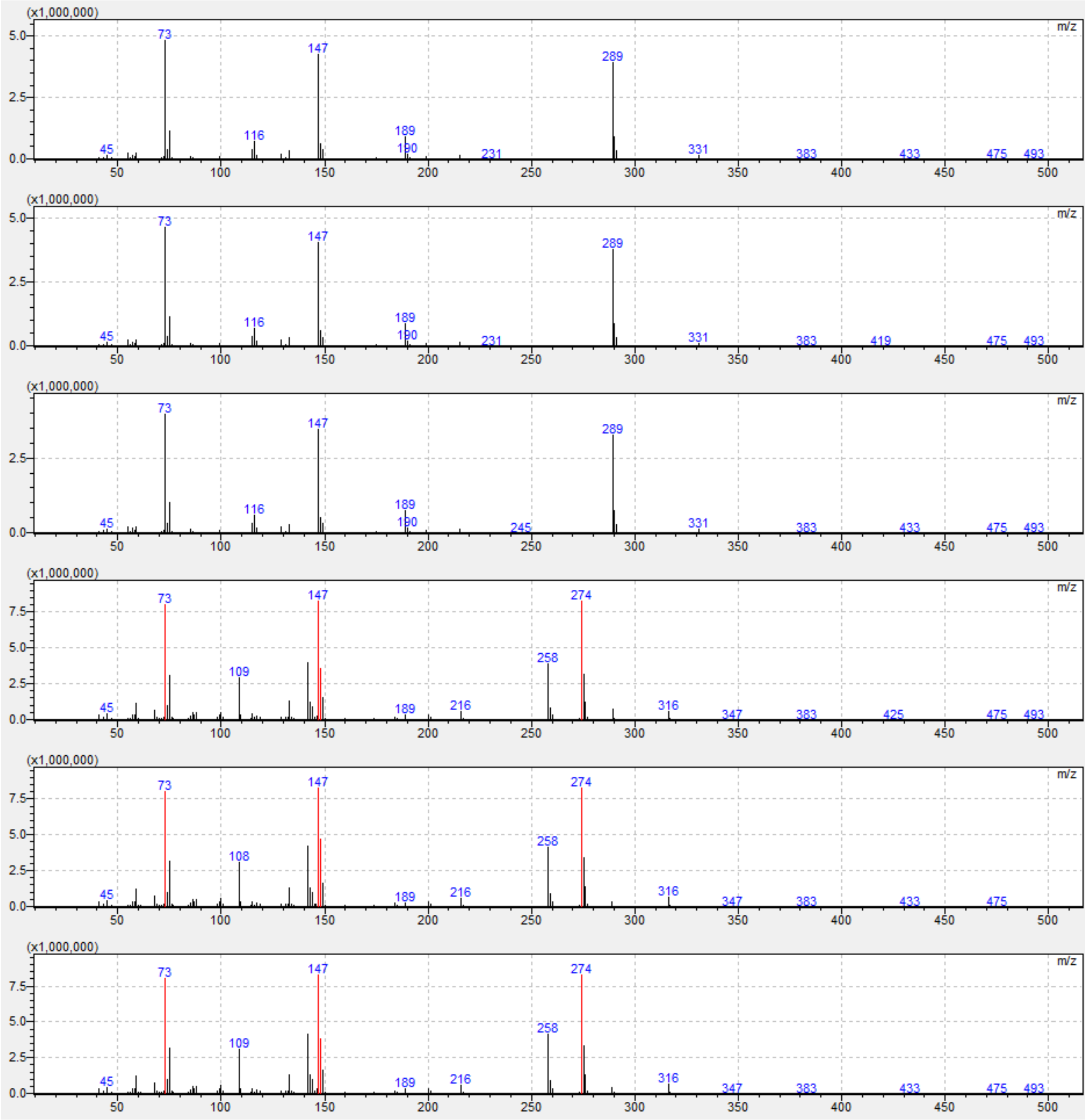


**Fig. S6.**

Representative GC-MS mass spectra at the retention time of γ-aminobutyric acid. Spectra from strain ASA549 (top three panels, three biological replicates) and strain ASA557-6 (bottom three panels, three biological replicates) are shown. This figure supports the data presented in Fig. 3A, B and Fig. S4.

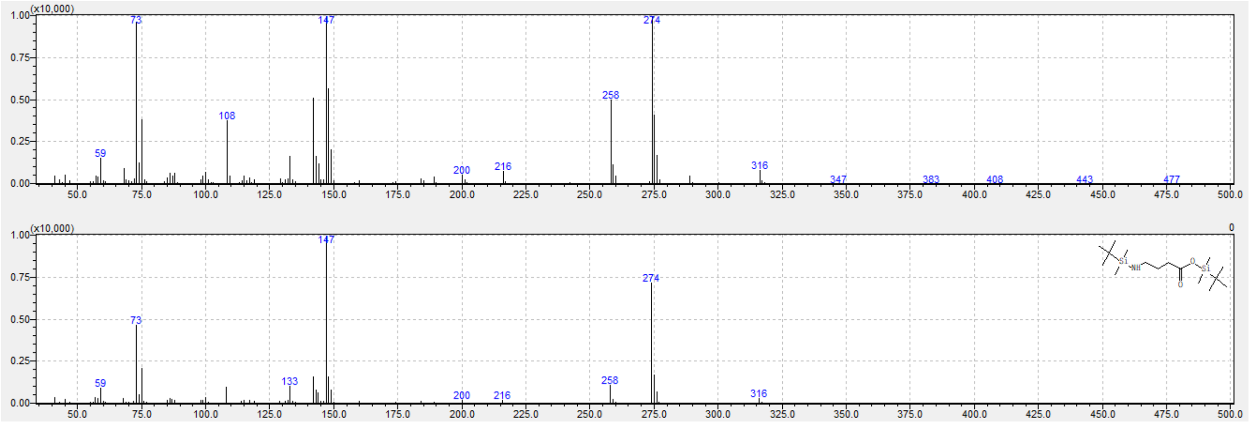


**Fig. S7.**

Comparison of a representative mass spectrum from strain ASA549 (top) with the γ-aminobutyric acid (GABA) reference spectrum from the library (bottom). This figure supports the data presented in Fig. 3A, B and Fig. S4.

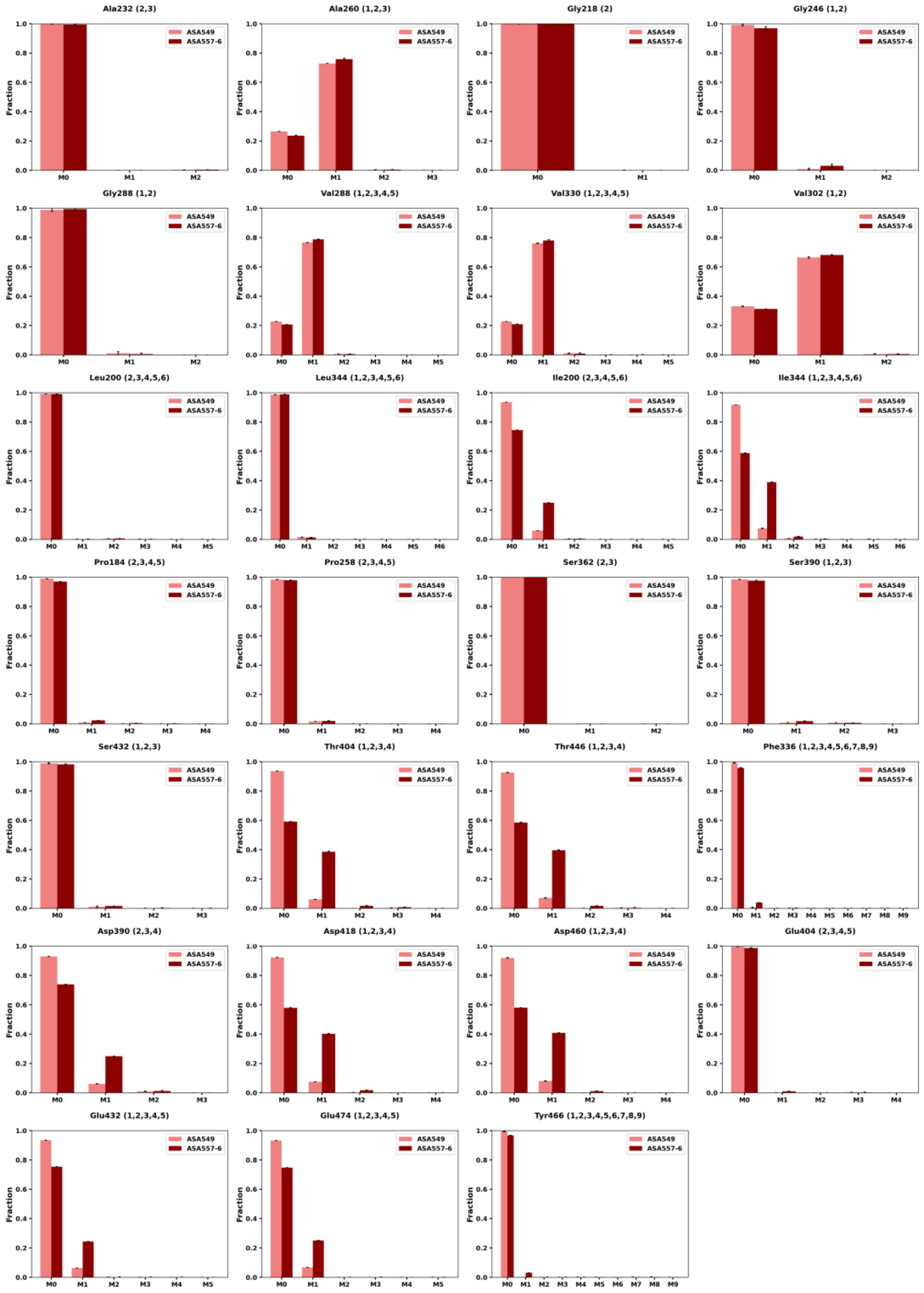


**Fig. S8.**

Mass isotopomer distribution of proteinogenic amino acid fragments from ASA549 and ASA557-6 grown on [1-$^{13}$C] D-glucose. Numbers in parentheses denote the carbon positions covered by each fragment. Data represent mean values and error bars indicate standard deviations from three technical replicates.

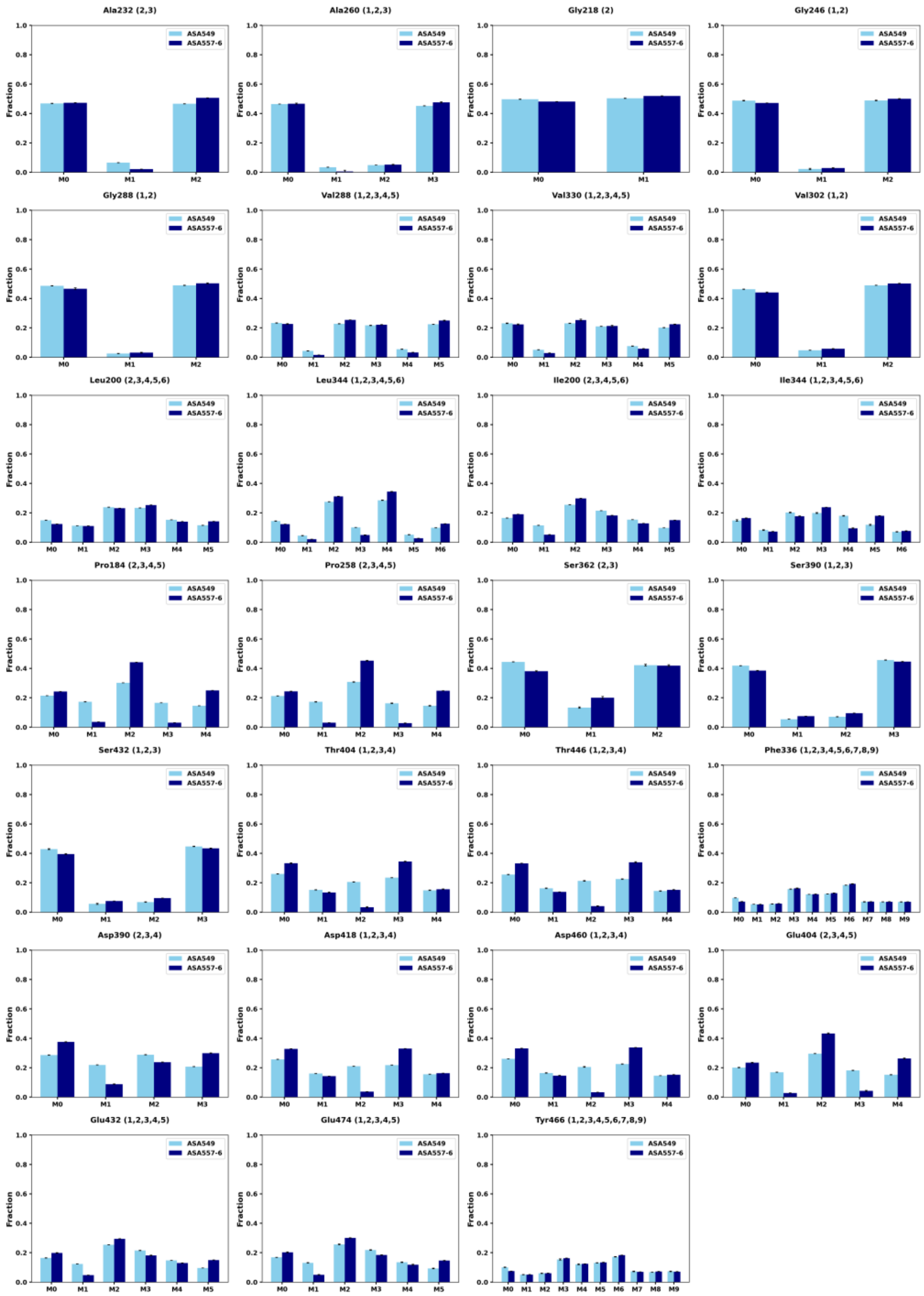

**Fig. S9.**

Mass isotopomer distribution of proteinogenic amino acid fragments from ASA549 and ASA557-6 grown on 50% [U-$^{13}$C] D-glucose. Numbers in parentheses denote the carbon positions covered by each fragment. Data represent mean values and error bars indicate standard deviations from three technical replicates.

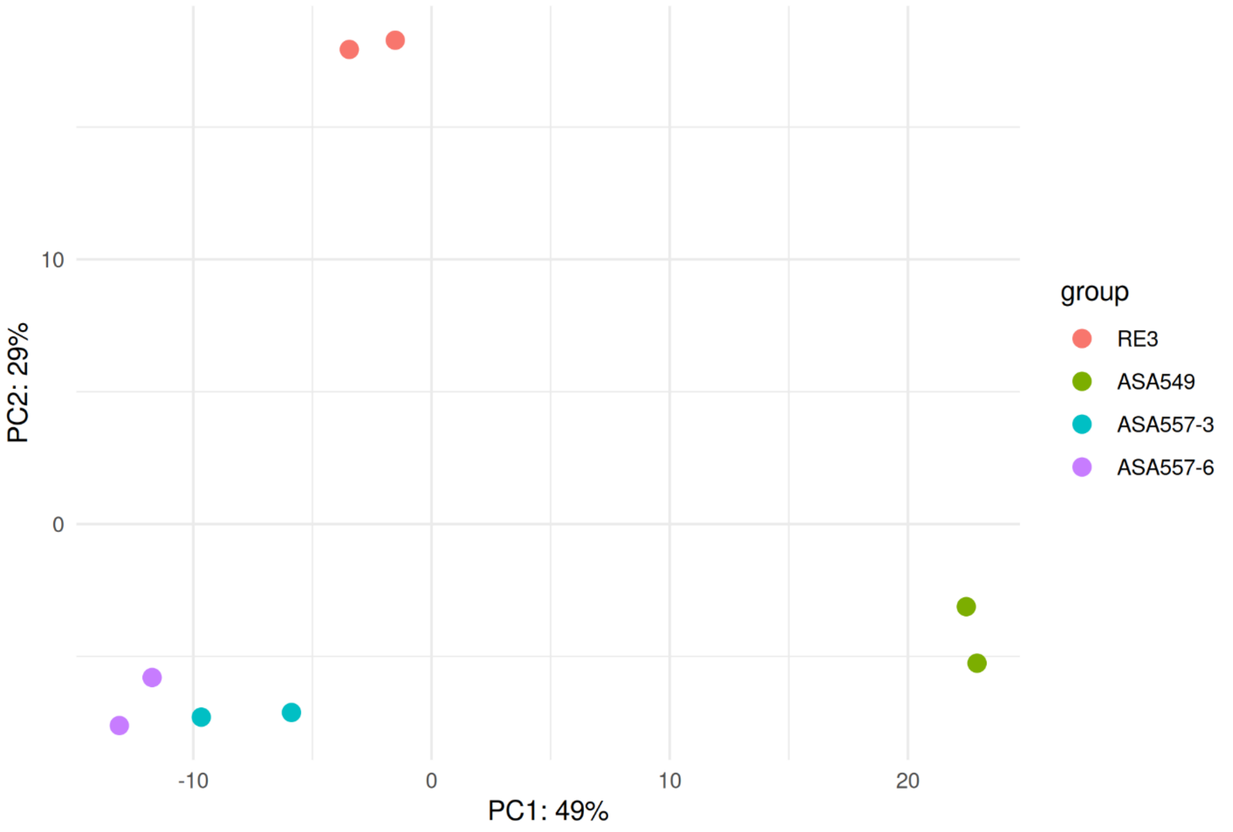


**Fig. S10**

Principal component analysis of top 500 differentially expressed genes of the ALE isolates ASA557-3 and ASA557-6 and reverse engineered strain RE3 with parental strain ASA549.

**Table S1.**

Standard curves for different fragments of metabolites analyzed by GC–MS.

| Fragment | Std. 1 (nmol) | Std. 2 (nmol) | Std. 3 (nmol) | Std. 4 (nmol) | Std. 5 (nmol) | Standard curve | R-squared |
|---|---|---|---|---|---|---|---|
| Ala158 | 10 | 40 | 80 | NA | NA | y = 2E+11x - 403219 | 0.9943 |
| Ala232 | 10 | 40 | 80 | NA | NA | y = 1E+11x - 197067 | 0.9953 |
| Ala260 | 10 | 40 | 80 | NA | NA | y = 1E+11x - 224292 | 0.9939 |
| Gly218 | 10 | 40 | 80 | NA | NA | y = 1E+11x - 165757 | 0.9938 |
| Gly246 | 10 | 40 | 80 | NA | NA | y = 1E+11x - 212085 | 0.9919 |
| Gly288 | 10 | 40 | 80 | 200 | 400 | y = 4E+09x + 2028.4 | 0.9955 |
| Leu200 | 1 | 5 | 10 | 20 | 50 | y = 3E+11x - 222385 | 0.9982 |
| Leu274 | 1 | 5 | 10 | 20 | 50 | y = 1E+11x - 60640 | 0.998 |
| Leu344 | 1 | 5 | 10 | 20 | 50 | y = 6E+09x - 4243.1 | 0.9977 |
| Ile200 | 1 | 5 | 10 | 20 | 50 | y = 3E+11x - 160131 | 0.9986 |
| Ile274 | 1 | 5 | 10 | 20 | 50 | y = 1E+11x - 36060 | 0.9983 |
| Ile344 | 1 | 5 | 10 | 20 | 50 | y = 5E+09x - 3356.9 | 0.9986 |
| Suc289 | 1 | 5 | 10 | 20 | 50 | y = 3E+11x - 73851 | 0.9985 |
| Pro258 | 1 | 40 | 80 | 200 | NA | y = 8E+10x - 490885 | 0.9955 |
| Pro286 | 1 | 40 | 80 | 200 | NA | y = 7E+10x - 571493 | 0.9911 |
| Fum287 | 0.1 | 5 | 10 | NA | NA | y = 4E+11x - 6208.1 | 1 |
| Met218 | 0.1 | 5 | 10 | 20 | 50 | y = 3E+11x - 582191 | 0.9915 |
| Met292 | 0.1 | 5 | 10 | 20 | 50 | y = 3E+11x - 512984 | 0.9905 |
| Met320 | 0.1 | 5 | 10 | 20 | 50 | y = 2E+11x - 415761 | 0.9915 |
| Ser432 | 1 | 40 | 80 | 200 | NA | y = 1E+10x - 22146 | 0.9996 |
| Ser302 | 1 | 40 | 80 | 200 | NA | y = 8E+10x - 126515 | 0.9997 |
| AKG346 | 1 | 5 | 10 | 20 | 50 | y = 2E+11x - 193131 | 0.9977 |
| Thr376 | 1 | 40 | 80 | 200 | 400 | y = 8E+09x - 30371 | 0.9993 |
| Thr404 | 1 | 40 | 80 | 200 | 400 | y = 1E+10x - 73914 | 0.9993 |
| Thr446 | 1 | 40 | 80 | 200 | 400 | y = 1E+09x - 6663.1 | 0.9992 |
| Phe234 | 1 | 5 | 10 | 20 | 50 | y = 2E+11x - 91111 | 0.9982 |
| Phe308 | 1 | 5 | 10 | 20 | 50 | y = 1E+11x - 31478 | 0.9977 |
| Phe336 | 1 | 5 | 10 | 20 | 50 | y = 1E+11x - 62351 | 0.9979 |
| Phe378 | 1 | 5 | 10 | 20 | 50 | y = 7E+09x - 4149.6 | 0.9979 |
| Mal391 | 1 | 5 | 10 | 20 | 50 | y = 3E+10x - 4867.3 | 0.9951 |
| Mal419 | 1 | 5 | 10 | 20 | 50 | y = 2E+11x - 84146 | 0.9967 |
| Asp316 | 10 | 40 | 80 | 200 | NA | y = 8E+10x - 132864 | 0.9998 |
| Asp390 | 10 | 40 | 80 | 200 | NA | y = 8E+10x + 109714 | 0.9995 |
| Asp460 | 10 | 40 | 80 | 200 | 400 | y = 5E+09x + 63199 | 0.9849 |
| Glu404 | 100 | 200 | 400 | NA | NA | y = 1E+10x + 499523 | 0.996 |
| Glu474 | 100 | 200 | 400 | 1000 | NA | y = 4E+09x + 459033 | 0.9881 |
| Cit431 | 0.1 | 5 | 10 | 20 | 50 | y = 1E+11x - 70669 | 0.9991 |
| Cit459 | 0.1 | 5 | 10 | 20 | 50 | y = 3E+11x - 235835 | 0.9989 |
| Tyr364 | 0.1 | 5 | 10 | 20 | 50 | y = 5E+10x - 11546 | 0.9987 |
| Tyr438 | 0.1 | 5 | 10 | 20 | 50 | y = 5E+10x - 3228.4 | 0.9987 |
| Tyr466 | 0.1 | 5 | 10 | 20 | 50 | y = 1E+11x - 28279 | 0.9988 |
| Tyr508 | 0.1 | 5 | 10 | 20 | 50 | y = 1E+10x - 4574.7 | 0.9987 |
| Tyr302 | 0.1 | 5 | 10 | 20 | NA | y = 7E+11x - 95147 | 0.9911 |

* Standard curves of the shaded fragments were used for the quantification of the corresponding metabolites.
